# Myosin 2 drives actin contractility in fast-crawling species outside of the amorphean lineage

**DOI:** 10.1101/2025.05.16.654244

**Authors:** Sarah L. Guest, Katrina B. Velle, Samantha M. Jacques, Yvette Park, Jarrett Man, Margaret A. Titus, Lillian K. Fritz-Laylin

## Abstract

Myosin 2-dependent actin contractility drives essential cell functions including fast crawling motility in animal cells, *Dictyostelium* amoebae, and other species from the Amorphea lineage. Whether and how species outside this single eukaryotic group can generate contractile actin networks has been largely unexplored. We demonstrate that *Naegleria,* an amoeba from the Heterolobosea—an evolutionarily distant eukaryotic lineage that includes the fastest known crawling eukaryotes—expresses three distinct Myosin 2 homologs. Using biochemical assays and immunofluorescence, we show that these Myosin 2 proteins bind cellular actin networks and that these networks generate ATP-dependent contractility. By identifying additional Myosin 2 homologs in dozens of additional heterolobosean amoebae (but not obligate flagellates), we find a widespread correlation within this group between crawling behavior and contractile actin networks. This correlation includes the amoeba *Vahlkampfia avara*, which we demonstrate can crawl at speeds exceeding 180 μm/min and has contractile actin networks and Myosin 2 homologs. These findings show that Myosin 2-driven contractility exists beyond Amorphea and is associated with diverse, fast-crawling cell types. Expanding the taxonomic breadth of actin network contractility impacts our basic understanding of cell motility, evolutionary biology, and of the fundamental biology of human pathogens that rely on fast cell migration.

## INTRODUCTION

Species from across the eukaryotic tree use forces generated by actin polymer networks to drive a wide range of cell functions. Human leukocytes and *Dictyostelium* amoebae, for example, use actin networks to rapidly migrate while hunting bacterial prey. This form of cell motility relies on cell shape changes that are, in part, driven by the contraction of actin polymer networks by Myosin 2 motor proteins.^1–3^ These motors form bipolar filaments that can pull on multiple actin filaments, sliding them past each other to contract the network.^4,5^ As the only myosin known to induce actin network contraction, Myosin 2 plays a crucial role in actin-dependent cell crawling in humans and other species. Beyond motility, contractile actin networks in these cells also drive other important functions, including cytokinesis, cell adhesion, and muscle contraction. Despite its clear importance to the cell types that have it, Myosin 2 activity has only been described within the Amorphea^6^—a single group of eukaryotes that includes animals, fungi, and *Dictyostelium* amoebae^7^ (**Figure 1A**, left)—raising questions about whether and how the vast majority of eukaryotes use contractile actin networks for crawling motility and other functions.

**Figure 1:**
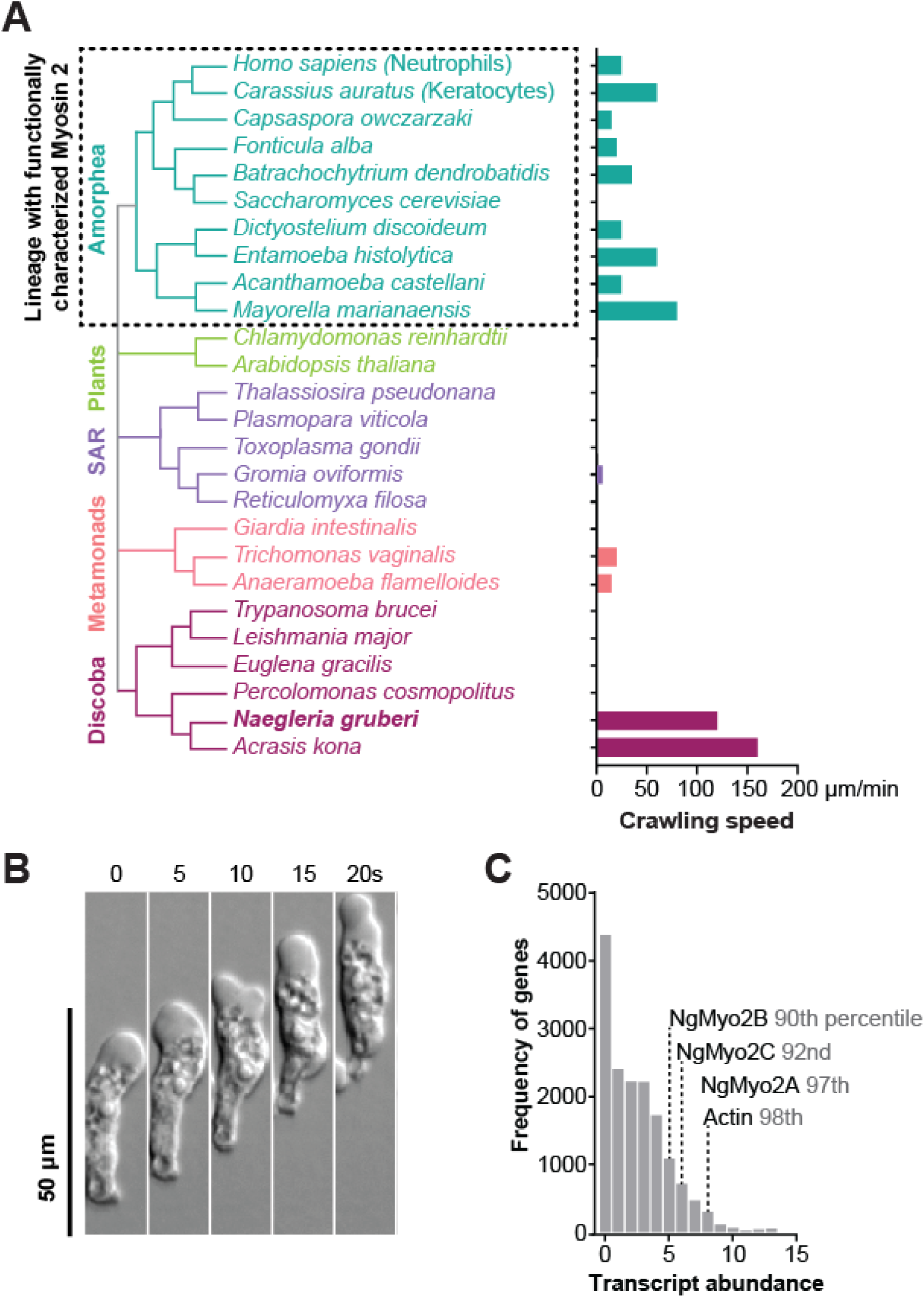
Myosin 2 exists outside of Amorphea, in the fast-crawling amoeba *Naegleria gruberi*. (**A**) A simplified species tree of representative eukaryotes from major lineages, and their maximum reported cell crawling speeds (bar graph, right).^75–85^. The Amorphea (teal) are the only lineage with functionally characterized Myosin 2. *Naegleria gruberi* and *Acrasis kona* are heterolobosean amoebae from the Discoba lineage (maroon) with top crawling speeds of 120 and 160 μm/minute.^79^ The species without values are not capable of crawling motility or lack published data on speed (*Reticulomyxa filosa*). (**B**) *Naegleria* amoebae crawl quickly across surfaces. A montage of a representative amoeba crawling on glass, imaged over 20 seconds using DIC. (**C**) *Naegleria* amoebae express three Myosin 2 genes. A histogram shows relative transcript abundance of genes expressed in amoebae. Frequency of values is shown for each bin of calculated abundance, calculated as log2-transformed average transcripts per million (TPM+1) for 5 replicates. The bins containing NgMyo2s (Myo2A, Myo2B, Myo2C) and the major actin gene (jgi_56107) are noted.

Within the Amorphea lineage, Myosin 2-dependent contraction of actin networks is central to the motility of many cells, including those that crawl with speeds greater than ten μm/min (**Figure 1A**, right).^8,9^ Fish keratocytes, for example, rely on actin-filled protrusions to push out their fronts and use Myosin 2 to contract the cell rear, enabling crawling speeds up to 60 μm/minute.^10,11^ Myosin 2 can also contribute to another mode of fast cell movement called blebbing motility by contracting the actin cortex—a meshwork of actin filaments beneath the plasma membrane—to form blister-like plasma membrane protrusions called blebs. These blebs can drive movement in animal cells such as primordial germ cells that crawl at 2.5 μm/min and amoebae like *Dictyostelium* whose speed can exceed 20 μm/min.^12,13^ Moreover, Myosin 2 inhibition by gene disruption or by drug perturbation reduces the speed of cells crawling using blebbing motility, as well as cells that use actin-filled protrusions.^14,15^ Despite the dependence of fast cell crawling on Myosin 2, its activity has only been characterized in Amorphea, leading to a general assumption that Myosin 2 was an amorphean innovation.^16^ However, the fastest known crawling cells are not from Amorphea, but instead are from an evolutionarily distant lineage called Heterolobosea^17^ that diverged over a billion years ago.^18^ This group includes the “brain-eating” amoeba *Naegleria fowleri*, and the related nonpathogenic model system *Naegleria gruberi,* whose crawling speeds can exceed 100 μm/minute (**Figure 1B**).^19^ The mechanisms driving the fast cell migration of these organisms, or any heterolobosean species, is severely understudied.

Recent work has revealed that *Naegleria* amoebae crawl using dynamic actin networks.^20^ Although its primary mode of crawling appears to rely on actin-filled protrusions called pseudopods,^20^ *Naegleria* can also crawl using blebs.^21^ The ability to bleb hints at a yet unidentified mechanism for actin cortex contraction, an activity that has only been attributed to Myosin 2. Accordingly, genome sequencing of *Naegleria gruberi* unearthed not one but three Myosin 2 genes (Myo2A, Myo2B, Myo2C),^6,22,23^ whose predicted motor domains are 58%-69% identical to each other and 47-51% identical to those of *Dictyostelium*. The discovery of Myosin 2 homologs in an evolutionarily distant Heterolobosean challenges the assumption that Myosin 2 activity is restricted to the amorphean lineage, but left more questions than answers about the origin and function of contractile actin networks in species outside the Amorphea.

Here, we use a combination of biochemical and cellular assays to show that *Naegleria* amoebae express all three Myosin 2 genes and that their proteins bind to and contract actin polymer networks. We also use fast crawling as a predictor of Myosin 2 content and contractile function in a previously unsequenced amoeba, and provide evidence that Myosin 2 homologs are widespread throughout the heterolobosean lineage. This work demonstrates that Myosin 2-mediated contractility is used outside of the Amorphea, and is a hallmark of fast crawling motility.

## RESULTS

### *Naegleria* amoebae express three Myosin 2 homologs

To determine if and how Myosin 2-dependent actin network contractility is used by species outside of the Amorphea lineage, we first determined whether and when *Naegleria* Myosin 2 genes are expressed. We quantified the relative transcript abundance of all three Myosin 2 genes using published RNA-seq data from actively growing cultures^24^ and found that all three Myosin 2s were highly expressed **(Figure 1C**). To determine whether Myosin 2 expression profiles vary over the cell cycle, we measured changes in transcript abundance in a published timecourse of synchronously dividing cells.^24^ Using mitotic tubulin expression as an indicator for mitosis,^24,25^ we compared the changes in Myosin 2 transcript abundance across the cell cycle. Average Myosin 2 transcript abundance peaks at 40 minutes after mitotic induction (**Figure 2A, top**), coinciding with the expected timing of cell division. Because Myosin 2 plays a key role in force generation during amorphean cytokinesis by contracting cortical actin networks, similar to its role in blebbing motility, the observed increase in *Naegleria* Myosin 2 expression during division reflects a potentially conserved role in both activities.^26^ Taken together, these data indicate that *Naegleria* amoebae actively transcribe all three Myosin 2s and suggest that Myosin 2s could play key roles in contractile force generation.

**Figure 2:**
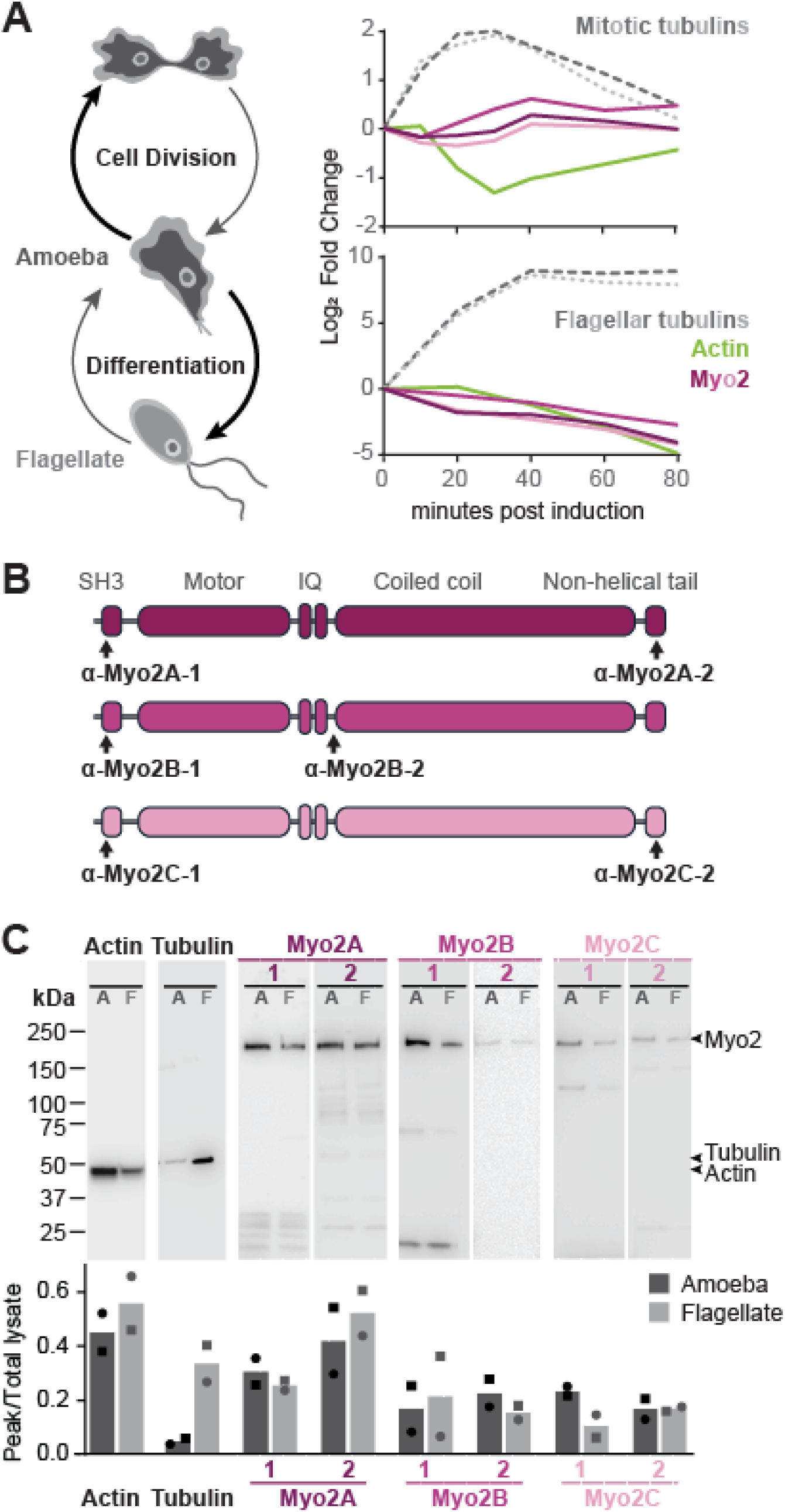
*Naegleria* expresses all three Myosin 2 genes. (**A**) *Naegleria* undergoes mitosis as an amoeba, but transiently differentiates into a swimming flagellate stage under stress (diagram, left). Expression of Myosin 2 varies during mitosis (top graph) and differentiation (bottom graph), shown as an adjusted log2 fold-change of each gene over 80 minutes, relative to 0 minutes post-induction. Cytokinesis occurs at 40 minutes after mitotic induction. The lines are color coded according to the genes expressed; green = actin, dark pink = Myo2A, pink = Myo2B, light pink = Myo2C. Mitotic tubulins (top graph, dark gray α-tubulin = AKG_0016; light gray β-tubulin = jgi_53354) are shown as a reference for mitotic progression. Flagellar tubulins (bottom graph, dark gray α-tubulin = jgi_39221; light gray β-tubulin = AKG_0052) are shown as an indicator for differentiation. (**B**) A cartoon of the domain architecture of *Naegleria’s* Myosin 2s indicates where the custom polyclonal antibodies bind; black arrows mark names and sites of peptide sequences used. Myo2A = dark pink, Myo2B = medium pink, Myo2C = light pink. (**C**) *Naegleria’s* Myosin 2s are expressed at the protein level. Western blot validation of antibodies raised against each Myosin 2 (top panel), and the relative quantitation (bottom panel) of proteins between amoebae (“A”, dark gray) and flagellates (“F”, light gray) for two replicates. Entire lanes are shown for assessing Myo2 degradation. Values were normalized between lanes of each replicate by total lysate quantitation.

In addition to growing and dividing as an amoeba, when stressed, *Naegleria* can transiently exit the cell cycle and differentiate into a swimming flagellate. During this amoeba-to-flagellate differentiation, *Naegleria* stops crawling and builds two flagella that it then uses to swim.^25,27,28^ Previous work has shown that *Naegleria* differentiation involves downregulation of genes encoding actin and upregulation of genes encoding flagellar tubulins.^24,29^ To explore the potential role of *Naegleria* Myosin 2 homologs in differentiation, we compared their transcript abundance to those of actin and flagellar tubulin during differentiation. While flagellar tubulin expression increased during differentiation, the expression of actin and all three Myosin 2s decreased (**Figure 2A, bottom**), consistent with a role for Myosin 2 in *Naegleria* amoebae rather than flagellates.

Having established that *Naegleria* expresses Myosin 2 at the RNA level, we next confirmed expression at the protein level. To this end, we generated custom polyclonal antibodies raised to both the N- and C- terminus of each Myosin 2. We selected N-terminal target epitopes near the Myosin 2 SH3 domain and C-terminal epitopes in the non-helical tailpiece (**Figure 2B**) and confirmed that the purified antibodies detected proteins of the expected molecular weights of each Myosin 2 in western blots of whole-cell extracts of *Naegleria* amoeba and flagellates (**Figure 2C**). In contrast to Myosin 2 mRNA levels that were reduced in flagellates compared to amoebae, we found no consistent change in Myosin 2 protein levels in extracts from both cell types. This expression pattern is similar to that of actin, whose mRNA abundance drops starkly during differentiation,^20^ but whose protein levels remain similar in both amoeba and flagellate.^30^ The actin in flagellates is thought to serve as a reservoir to facilitate the rapid return to crawling motility upon de-differentiation,^29^ hinting that Myosin 2 proteins may too. Taken together, these results show that *Naegleria* amoebae express all three Myosin 2 genes.

### Naegleria Myosin 2 proteins bind to the actin cortex

In Amorphea, Myosin 2 associates with the actin cortex, where it imparts contractile forces that drive cell crawling.^9,11^ To explore whether *Naegleria* Myosin 2 may function similarly, we tested whether *Naegleria* Myosin 2 proteins localize to the actin cortex by immunofluorescence. All three N-terminal antibodies localized to punctate structures that appear concentrated at the cell periphery, overlapping with actin (**Figure 3A**). To quantify this impression, we used line scans to measure myosin and actin fluorescence intensities across the cell, passing through both sides of the cell cortex. This analysis revealed overlapping peaks of actin and Myosin 2 signal at the cell periphery, as well as within the cell interior. To assess the overlap of Myosin 2 with actin inside the cell, we used a 90° ROI rotation colocalization test (see **Methods, Dataset S1**).^31^ The average Pearson’s correlation coefficient of actin:Myosin 2 colocalization across all three antibodies was weakly positive; (0.30 ± 0.04; average ± SE), while the control measurements of 90°:0° rotated images had scores close to zero (-0.05 ± 0.05; average ± SE). Because tubulin does not typically colocalize with Myosin 2, we performed the same colocalization test using tubulin:actin signal correlation as a control in both amoebae and flagellates (**Figure S1**). Both Pearson’s correlations were close to zero, indicating no relationship between localization of actin and tubulin (-0.06 for flagellates and 0.11 for amoebae). The lack of measurable correlation between tubulin and actin validates the correlation between actin and Myosin 2 at the periphery, as well as the less obvious co-localization in the cell interior. These measurements indicate a biased localization of all three Myosin 2 proteins to cortical actin networks.

**Figure 3:**
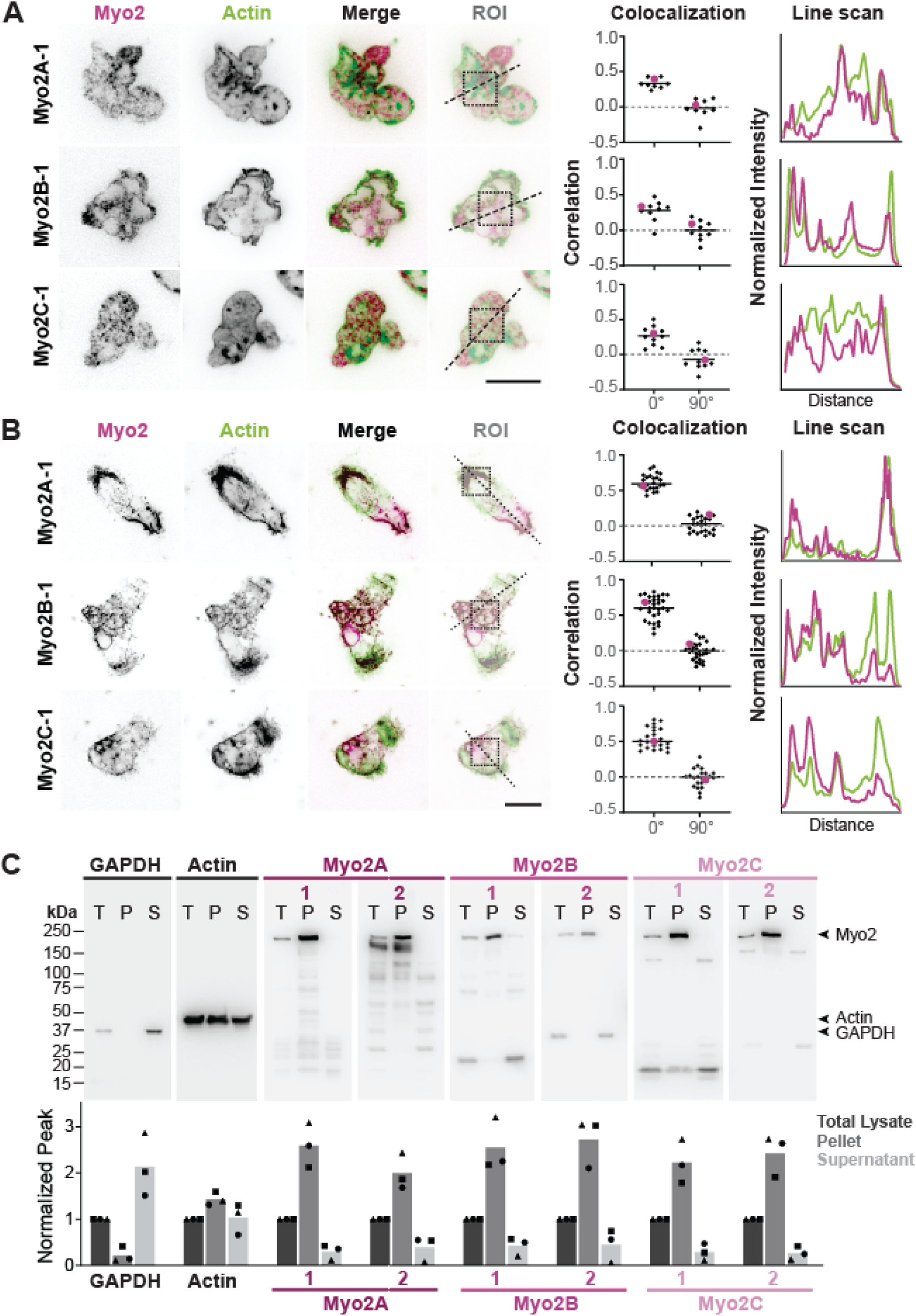
*Naegleria’s* Myosin 2s bind actin networks. (**A**) *Naegleria* amoebae were fixed and stained with Myo2 antibodies (pink) to detect Myosin 2 and Alexa Fluor™ 488 Phalloidin to label actin filaments (green). Single z slices through the midsection of cells are shown. The Region of Interest (ROI) for each example depicts the area used for colocalization analyses; dotted squares were used in 90° rotation tests, and dotted lines through the peripheral actin cortex were used for line scans (far right). The colocalization values are plotted as Pearson’s correlation coefficients, with the representative cells shown on the left indicated by pink circles. The Myo2 and actin line scan fluorescence intensity was normalized to the dimmest (0%) and brightest (100%) signal for regions within the cortex boundary. (n ≥ 6 cells for each antibody; scale bar = 10μm). (**B**) Myosin 2 remains bound to *Naegleria’s* extracted actin cytoskeleton (ghosts). Following cytoskeletal extraction (see Methods), ghosts were fixed, stained and analyzed as in (A). (n ≥ 21 ghosts per antibody; scale bar = 10μm). (**C**) Myosin 2 is retained in the insoluble fraction of protein lysates. Cells were lysed and protein samples were separated into cytoskeletal pellet and soluble fractions, then subjected to western blot analysis with all 6 Myosin 2 antibodies. Actin and GAPDH antibodies were used as controls for cytoskeletal pellet and soluble supernatant, respectively. Quantitation of three replicates shows peak intensity normalized to total lysate per lane for each replicate trio. T= Total Lysate (dark gray), P = Pellet (medium gray), S = Supernatant (light gray).

Given the intense staining of each antibody within the cell body, we hypothesized that a large fraction of *Naegleria* myosin could be present in an unbound, cytosolic pool. To specifically visualize the actin-bound fraction of Myosin 2, we extracted the cells’ cytoskeletons by lysing with detergent in the presence of fluorescent phalloidin to stabilize actin polymers, washed to remove cytosol and unbound cell contents, then immunostained with anti-Myosin 2 antibodies.^32^ These extracted cytoskeletons, called “ghosts”, remained adhered to the surface while cytoplasm, organelles and other particles dispersed after lysis. Like in whole-cell immunofluorescence, N-terminal antibodies localized to the actin networks of *Naegleria* ghosts (**Figure 3B**). The ghosts, however, appeared to lack the diffuse cytoplasmic signal seen in whole-cell immunofluorescence. We repeated the ROI 90° rotation colocalization test and we found a strong correlation between actin and myosin in *Naegleria* ghosts (0:0° = 0.59 ± 0.03, vs 90°:0° = 0.00 ± 0.02). The increase in the correlation in ghosts compared to whole cell immunofluorescence is consistent with all three *Naegleria* Myosin 2s existing in both actin-bound and unbound cytosolic states.

We next quantified the proportion of Myosin 2 bound to actin by centrifuging cell lysates to separate the cortical actin cytoskeleton and associated proteins from soluble proteins and measured the relative amounts of insoluble (actin-bound) and soluble (free) Myosin 2 by Western blot (**Figure 3C**). We used a custom anti-Ng-GAPDH antibody as a positive control for the soluble fraction,^33^ and an anti-actin antibody as a positive control for the insoluble fraction.^34^ The amount of actin in the soluble fraction was similar to that in the insoluble fraction, consistent with a 1:1 ratio of polymerized to unpolymerized actin, as has been reported for *Dictyostelium* amoebae.^35^ In contrast, the Myosin 2 signal was approximately four times greater in the insoluble fraction, confirming that *Naegleria’s* Myosin 2s are physically associated primarily with polymerized actin. Taken together, these data show that *Naegleria* Myosin 2 proteins exist in two states: an unbound, cytosolic state, and an actin-bound state enriched at the cell cortex.

### *Naegleria* Myosin 2 protein provides actin network contractility

Satisfied that *Naegleria* Myosin 2s bind polymerized actin, we next explored whether they contribute to the contraction of cellular actin networks. Actin contractility can be quantified using “ghost reanimation assays”, in which detergent-extracted actin cytoskeletons are treated with ATP to activate Myosin 2 motor activity, resulting in shrinkage of the network.^32,36,37^ To test if *Naegleria’s* actin network is contractile, we performed a ghost reanimation assay of *Naegleria*, using *Dictyostelium* amoebae as a positive control.^32^ We imaged ghosts during Mg-ATP addition and observed immediate and robust contractions of ghosts of both amoebae; *Dictyostelium* ghosts lost an average of 48.5% ± 4.6% (SE) of their original footprints, while *Naegleria* ghosts lost an average of 42.4% ± 6.5% (**Figure 4A, Supplemental Movie S1**). These data demonstrate that *Naegleria’s* actin network is contractile, and that like actomyosin contractile networks from amorphean species, its contractility is ATP dependent.

**Figure 4:**
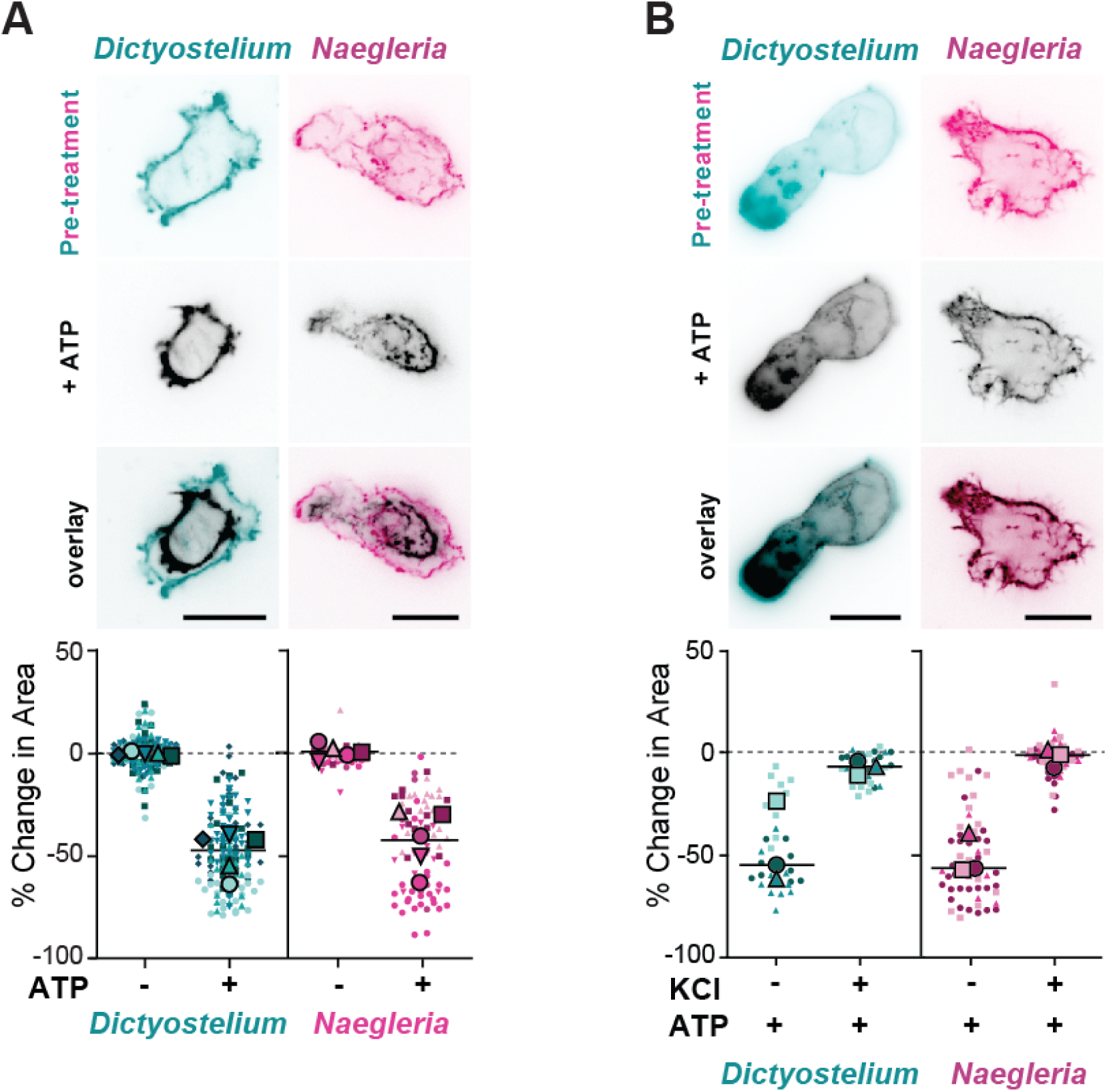
*Naegleria* actin networks are contractile in an ATP dependent manner. (**A**) *Naegleria’s* actin networks are contractile, like *Dictyostelium*. Images show representative *Dictyostelium* and *Naegleria* ghosts before (teal, pink) and after (black) the addition of 1mM Mg-ATP to activate Myosin 2. The graph shows ghost surface area contraction, calculated as the percent change in pixel area after addition of buffer (-) or 1mM Mg-ATP (+). (N = 5, scale bars = 10 μm). (**B**) Ghost contractility is abolished by salt concentrations that disrupt Myosin 2 filaments. *Dictyostelium* and *Naegleria* ghosts were treated with 200mM KCl and imaged before (teal, pink) or after the addition of ATP (black). Quantification was completed as in (A) for ghosts in wells with (+) and without (-) prior KCl treatment. (N = 3, scale bars = 10 μm)

A hallmark of Myosin 2-dependent actin network contraction is the loss of contractility upon salt treatment. This effect is caused by salt-induced disruption of electrostatic interactions mediating filament assembly, which leads to the breakdown of multimer filaments and loss of network contractility.^4,38–41^ To verify that *Naegleria*’s actin network contractility was dependent on Myosin 2 filaments, we treated cells with 200mM KCl at pH 7.6, which leads to loss of Myosin 2 activity in animal and *Dictyostelium* ghosts,^38,40^ then induced contraction by adding Mg-ATP while imaging (**Figure 4B**). We again used *Dictyostelium* as a positive control, and found that control ghosts contracted (46.6% ± 20.2% reduction), while salt-treated ghosts did not (7.2% ± 3.4% reduction). Similarly, *Naegleria* control ghosts showed robust contraction upon ATP addition (51.3% ± 4.8% reduction), while *Naegleria* ghosts treated with salt did not contract (2.3% ± 1.6% reduction). The loss of contraction upon salt treatment suggests that Myosin 2 filaments are likely to be important for driving *Naegleria’s* actin network contractility.

To further test whether *Naegleria* actin network contractility requires filamentous Myosin 2, we leveraged the ability of Myosin 2 filaments, but not monomers, to remain bound to actin through repeated ATP hydrolysis cycles.^42^ Under normal conditions, the motor domains of a large Myosin 2 filament do not simultaneously release actin, thereby maintaining contact with the actin network. In contrast, Myosin 2 monomers are free to diffuse away from the actin filament following ATP hydrolysis (**Figure 5A**).^32,38,43^ To test if *Naegleria’s* Myosin 2s dissociate from actin like those of Amorphea, we treated ghosts with KCl, Mg-ATP, or a double treatment with KCl followed by Mg-ATP. We then fixed and immunostained each set of treated ghosts with anti-Myosin 2 antibody and measured the total fluorescence per ghost **(Figure 5A)**. The salt-treated ghosts lost a substantial amount of Myosin 2 signal after the addition of Mg-ATP, but Myosin 2 was not lost in ghosts treated with KCl or ATP alone **(Figure 5B)**. The retention of Myosin 2 signal on ATP-treated ghosts indicates that ATPase activation is insufficient for Myosin 2 to release the actin network, consistent with its assembly into filaments. These results indicate that *Naegleria*’s Myosin 2s form multimers that are activated by ATP and sensitive to high salt, a characteristic of higher-order Myosin 2 filaments found in amorphean species. Collectively, these data support a model in which *Naegleria’s* Myosin 2s, like conventional amorphean Myosin 2s, are recruited to the actin cytoskeleton, form higher order filaments and induce actin network contraction in an ATP-dependent manner.

**Figure 5:**
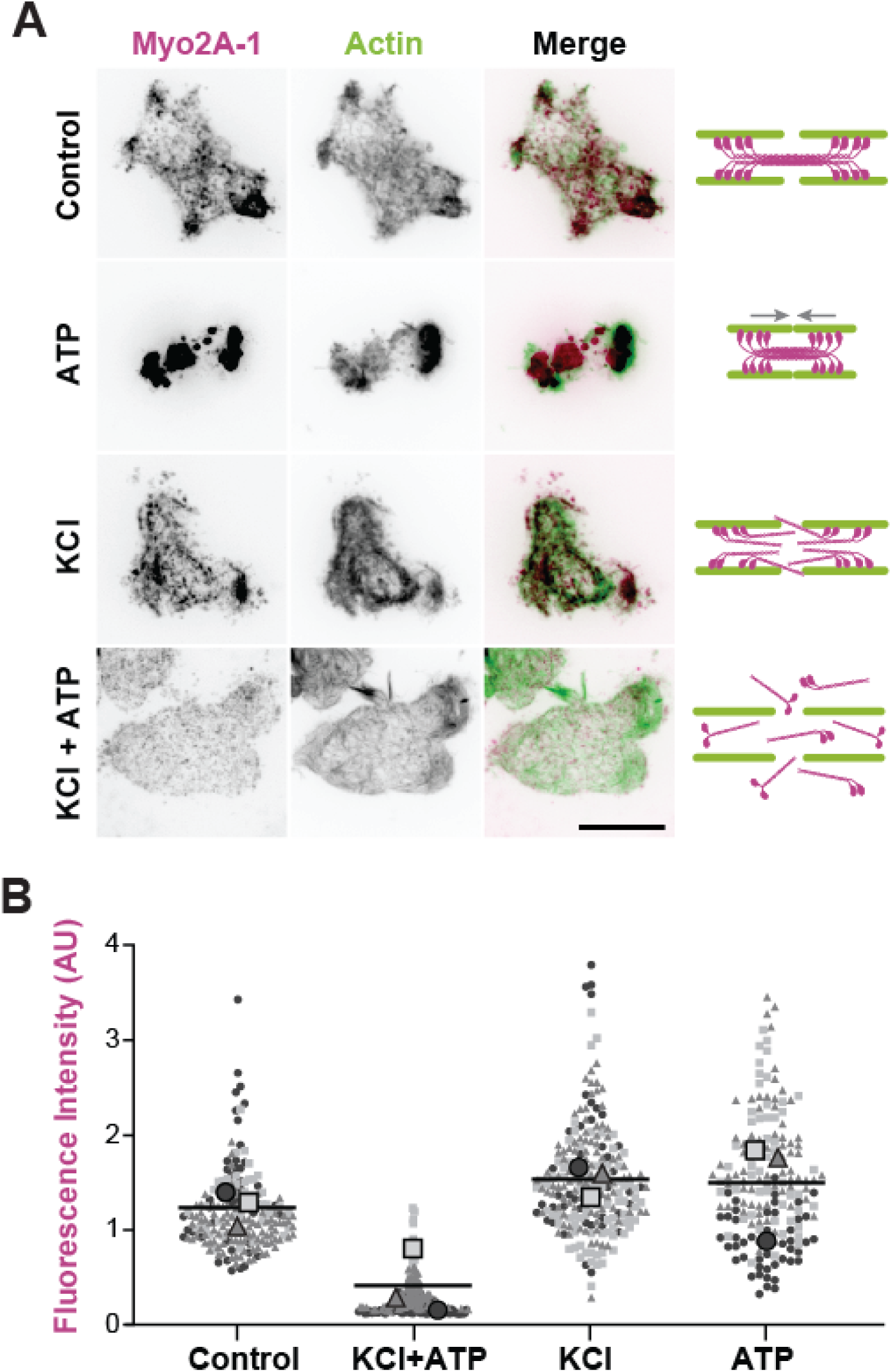
*Naegleria’s* Myosin 2s release actin under high salt with ATP. (**A**) Right: A schematic representation of the expected states of Myo2 filaments in each experimental condition. Left: Images of ghosts treated in the conditions as shown above with buffer alone (control), or buffer supplemented with: 200mM KCl (‘KCl’), 1mM Mg-ATP (‘ATP’), or sequentially with KCl 10 minutes, then Mg-ATP for 10 minutes (‘KCl+ATP’). The ghosts were then fixed and immunostained with Myo2A-1 antibody (pink) and labeled with phalloidin-488 (green). Scale bar = 10 μm. (**B**) Average fluorescence intensity quantified per ghost from maximum intensity-projections across 3 replicates.

### Other heterolobosean amoebae also have Myosin 2 homologs and contractile actin networks

Having shown that *Naegleria* Myosin 2 functions in actin contractility, we wondered whether other species outside of Amorphea might also use Myosin 2 for actomyosin contraction.

Previous searches found Myosin 2 only in Amorphea and two heterolobosean species: *Naegleria gruberi* and *Naegleria fowleri.*^6,23^ We searched for Myosin 2 in recently-published genomic and transcriptomic datasets from species spanning the eukaryotic tree. This search recovered putative Myosin 2 genes from 10 additional heterolobosean species (**Figure S2,** *Naegleria lovaniensis, Willaertia magna, Tetramitus jugosus, Acrasis kona, Neovahlkampfia damariscottae, Pharyngomonas kirbyi, Percolomonas cosmopolitus, Soginia anisocystis, Heteramoeba clara,* and *Allovahlkampfia sp. BA* ^44–49^) suggesting that contractile actin networks may be widespread across this eukaryotic group. Intriguingly, of all the heterolobosean species we queried, only those capable of crawling contained Myosin 2.^50,51^ Furthermore, targeted searches for Myosin 2 in *Percolomonas*, a non-crawling heterolobosean flagellate, revealed no candidate genes that group with the Myosin 2 gene clade.^52^ The presence of Myosin 2 homologs only in crawling heterolobosean species points to a broader relationship between Myosin 2 gene content and cell crawling, and suggests that heterolobosean species that do not crawl may lose Myosin 2.

To determine whether these new Myosin 2 genes function similarly to those from well-studied amorphean species, we compared these new homologs to *Dictyostelium* Myosin 2, focusing on specific regions that confer distinct biochemical activities. Specifically, we checked for an N-terminal SH3 domain involved in mediating motor kinetics and essential light chain interactions,^53,54^ specific motifs within the motor domain that confer its ATPase and actin binding activity,^55,56^ and exactly two IQ motifs that interact with myosin light chains for regulation^57^ using a variety of sequence analysis tools (**Figure 6A**). While some regions were more varied (SH3, CM loop, actin-interacting loops 1-4, IQ motifs, **Figure S3, S4**), most functional motifs within the motor domain had high similarity scores to the *Dictyostelium* reference (purine-binding loop, P-loop, Switch I and II, relay, SH1 and SH2 helices, and converters, **Figure 6A**). These data indicate that, while the heterolobosean Myosin 2 sequences diverge in predicted protein-protein interacting regions, the core ATPase and force transduction regions remain conserved to those of Amorphea, consistent with our biochemical characterization of *Naegleria* Myosin 2 proteins.

**Figure 6:**
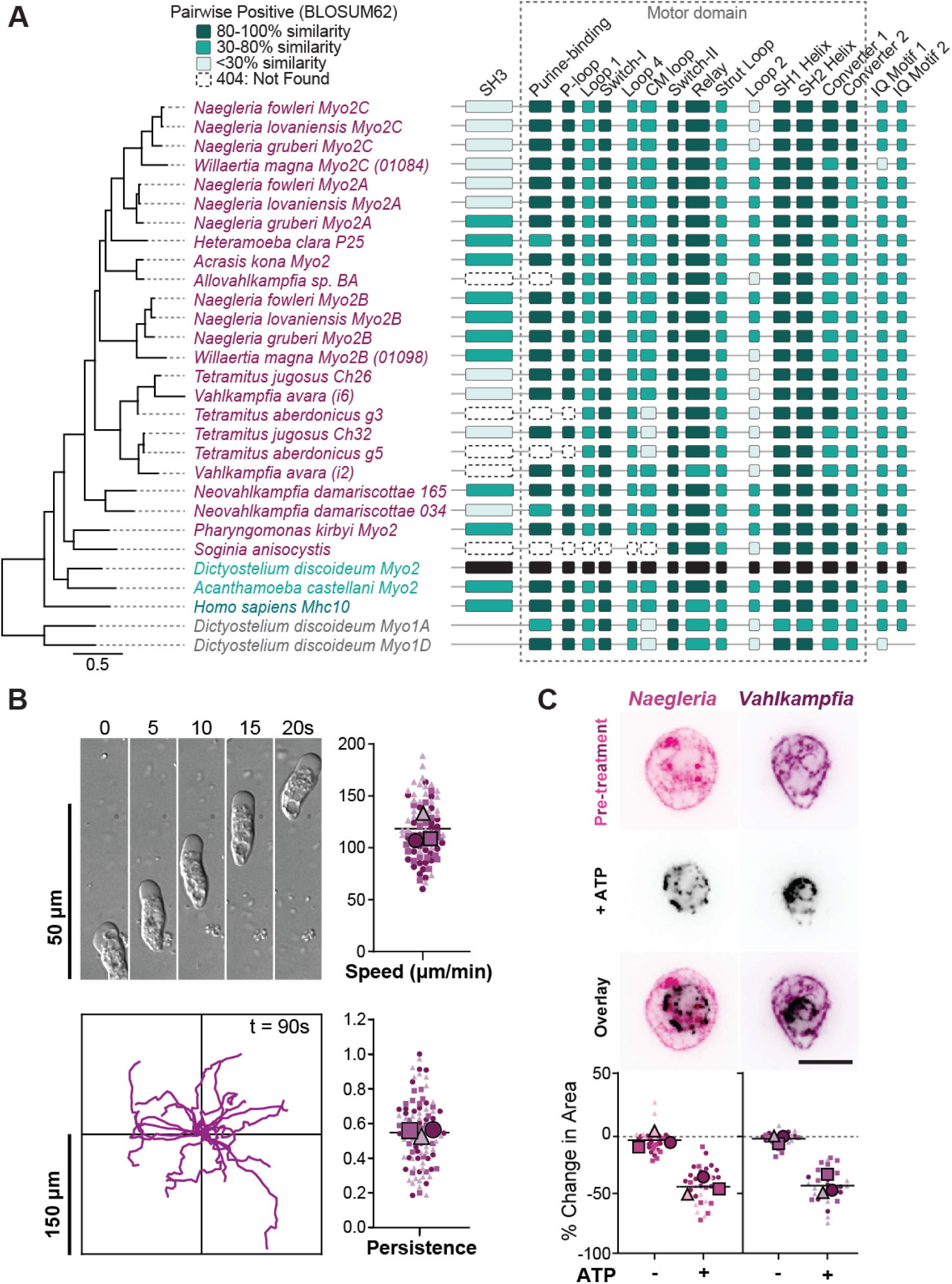
Other heterolobosean species also have Myosin 2. (**A**) A modified jelly bean plot depicts subdomain motifs in a multiple sequence alignment of heterolobosean Myosin 2s, aligned with ClustalO from the N-terminus through the second IQ motif, and organized phylogenetically with a tree built in Geneious using RAxML v 8.2.12, with a scale bar indicating substitutions per site. Highlighted subdomain motifs within (gray box) and proximal to the motor domain were colorized by pairwise positive scores via Blosum62 matrix, as compared to the *Dictyostelium* Myosin 2 sequence (shown in black). Scores above 80% similarity are shaded in dark teal, 30-80% in teal, and <30% in light teal. Missing regions of proteins due to partial or incomplete datasets are noted as “Not Found” (striped). (**B**) Another Heterolobosean*, Vahlkampfia avara,* also crawls quickly. A montage of typical *Vahlkampfia* amoebae crawling on an untreated glass surface (top left panel) shows morphology and representative speed. Individual cell tracks are shown for a 90-second timelapse (bottom left panel), normalized so time=0 is at the origin. Average crawling speed (top right panel) and directional persistence (maximum displacement divided by the total path length) are quantified (bottom right panel). N=3. (**C**) *Vahlkampfia* amoeba also have contractile actin networks. Images show representative *Naegleria and Vahlkampfia* ghosts before (pink, purple) and after (black) the addition of 1 mM Mg-ATP to activate Myosin 2. The graph shows ghost surface area contraction, calculated as the percent change in pixel area after addition of buffer (-) or 1 mM Mg-ATP (+). (N = 3, scale bars = 10 μm).

To explore the evolutionary history of Myosin 2 in Heterolobosea, we inferred a gene tree of all the putative heterolobosean Myosin 2 sequences, those of characterized amorphean Myosin 2s, as well as other myosin motors from diverse Eukaryotes as outgroups. The resulting gene tree topology supports a monophyletic clade of heterolobosean Myosin 2 (100% Bootstrap), and recapitulates the expected heterolobosean species tree within that branch,^58^ with gene duplications occurring prior to and throughout heterolobosean species divergence (**Figure S2)**. This topology is consistent with vertical inheritance of genuine Myosin 2 genes throughout Heterolobosea. Moreover, the presence of Myosin 2 genes along with crawling motility in many sequenced heterolobosean species raises the possibility that these two features may be tightly correlated.

Having established that Heterolobosea contains genuine Myosin 2 orthologs, we next sought to test the association between Myosin 2 and cell crawling by determining whether crawling can *predict* the presence of Myosin 2 genes. To this end, we investigated the presence of Myosin 2 in an unsequenced heterolobosean amoeba, *Vahlkampfia avara*. We cultured *Vahlkampfia* and confirmed that it could crawl. Surprisingly, these cells crawl on glass at an average of 97.4 μm/min, with fastest speeds over 180 μm/min, representing the fastest recorded crawling cell we are aware of **(Figure 6B)**. Moreover, *Vahlkampfia* amoebae have a directional persistence at 0.65 ± 0.03 across three replicates, a migration behavior similar to *Naegleria* (0.63 ± 0.04).^20^ To test whether *Vahlkampfia’s* actin networks are contractile, we again employed the ghost reanimation assay. We found that these cells also contract upon addition of Mg-ATP, with an average surface area reduction to 43.7% of original size, similar to that of *Naegleria* (**Figure 6C, Supplemental Movie S1**). This suggests that, like *Naegleria* and *Dictyostelium*, the cortical actin network of *Vahlkampfia* may use Myosin 2 for contractility. To test this hypothesis, we sequenced *Vahlkampfia* cDNA and found transcripts corresponding to multiple myosin genes. Phylogenetic analysis places two of these genes unambiguously in the Myosin 2 family (**Figure S2**), with motor domains 53-59% identical to *Naegleria* Myosin 2s. This indicates that fast cell crawling successfully predicted the presence of Myosin 2 in the *Vahlkampfia* genome.

## DISCUSSION

Despite the importance of contractile actin networks to the biology of animals, fungi, and their unicellular relatives, the existence of these networks in other major eukaryotic lineages was previously unknown. Our results demonstrate that the evolutionarily distant amoeba *Naegleria* expresses three Myosin 2 homologs that, like their amorphean counterparts, bind and contract actin networks. We also show that Myosin 2 genes are widespread in species throughout Heterolobosea, where fast crawling is common. We further demonstrate that the fast crawling motility of another Heterolobosean, *Vahlkampfia*, can predict the presence of both contractile actin networks and Myosin 2 genes. Taken together, these findings challenge the long-held assumption that Myosin 2–driven cortical contractility is confined to the amorphean lineage.

We previously showed that, like some amorphean cells, *Naegleria* can switch its crawling strategy between blebbing motility and actin-polymerization-dependent pseudopods.^20^ Based on what we know about the cell migration of animal and *Dictyostelium* cells, either of these strategies may rely on Myosin 2 activity.^11,14,15^ However, the precise role of Myosin 2 in either mode of migration in *Naegleria*, or any heterolobosean amoeba, awaits the development of robust genetic tools for these species. In addition to shedding light on the evolution of contractile actin networks, determining the mechanisms that drive fast cell migration in *Naegleria* also has direct relevance to human health, as this work could support the development of drugs targeting the deadly brain-eating amoeba *Naegleria fowleri,* whose infection pathway depends on its ability to rapidly migrate toward and within the brain.^59,60^

The similarity between heterolobosean and amorphean Myosin 2 functional domains (**Figure 6A**) suggests that their molecular mechanics are conserved across these deeply diverged lineages. This discovery raises an obvious question: Where did heterolobosean Myosin 2s come from? Previous work, based on gene trees that included only the *Naegleria* Myosin 2s, raised the possibility that these genes were horizontally transferred from an Amorphean species.^6^ Although our expanded Myosin 2 phylogeny neither supports nor undermines this hypothesis, either result is interesting. On the one hand, the discovery of Myosin 2 genes throughout Heterolobosea indicates that, if heterolobosean Myosin 2 originated via horizontal gene transfer, this event must have occurred over a billion years ago, prior to the diversification of this clade (1.1-1.8 Gya).^18^ If, on the other hand, heterolobosean Myosin 2 genes were vertically inherited, their presence within the heterolobosean lineage would suggest that this motor protein arose early in eukaryotic evolution—potentially even in the last common eukaryotic ancestor. This scenario would imply that Myosin 2, and the capacity to generate contractile actin networks, were lost independently in many eukaryotic lineages.

Regardless of whether heterolobosean Myosin 2 genes were acquired through vertical inheritance or horizontal gene transfer, their widespread retention across the clade points to a conserved and functionally significant role. Numerous studies in animal and *Dictyostelium* cells have highlighted the importance of Myosin 2 contractility to cell migration in Amorphea, where it is clearly important for high speed motility.^8,11,14,15^ Here, we posit that fast cell migration in the Heterolobosea also relies on Myosin 2-derived actin cortex contractility. This idea is supported by the correlation between crawling motility and the retention of Myosin 2 homologs within Heterolobosea, as well as our ability to predict the presence of Myosin 2 homologs in a non-sequenced species based on its crawling speed. The association of Myosin 2 with fast cell crawling in diverse eukaryotic groups raises the possibility that Myosin 2, and therefore contractile actin networks, may be present in other understudied or unsequenced lineages, prompting a reevaluation of our assumptions about the basic cell biology of diverse eukaryotes that are critical for human health, agriculture, and global ecosystems.

## ACKNOWLEDGEMENTS

We thank Ravi Ranjan for genomics service and sequencing, performed at Genomics Resource Laboratory, University of Massachusetts Amherst, MA (RRID:SCR_017907). We thank Dr. Marc Edwards for *Dictyostelium discoideum* cells. This work was primarily funded by NIH grant 1R35GM143039 to L.K.L.-F. And also supported by NIH grant R00GM147656 and a Research Award from Amazing Aven’s Quest for Amoeba Awareness to K.B.V., NIH grant R01GM122917 to M.A.T., and a Smith Family Foundation Award for Excellence in Biomedical Science to L.K.F.-L. who is Canadian Institute for Advanced Research (CIFAR) fellow in the Fungal Kingdom: Threats and Opportunities program and an Investigator of the Howard Hughes Medical Institute.

## METHODS

### Cell culture

*Naegleria gruberi* strain NEG-M (ATCC 30224) was grown axenically as previously described.^25^ Cells were grown to a density of ∼ 1 × 10^6^ cells/ml in M7 media (0.362 g/L KH2PO4, 0.5 g/L Na2HPO4, 5.4 g/L glucose, 5 g/L yeast extract (Difco), 45 mg/L L-methionine, 10% fetal bovine serum) at 28°C in tissue culture flasks. Cells were passaged every 2–3 days by ∼20-fold dilution, and maintained for no more than 30 passages before restarting from frozen stocks.

*Naegleria gruberi* strain NEG-M differentiation: Cells were grown to peak density then collected into a 15 ml conical and centrifuged at 900 x g for 5 minutes. The supernatant was removed, and cells were resuspended in 10 ml of pre-chilled 2 mM Tris pH 7.2. This was repeated for a total of three washes. The cells were then transferred into a sterile Erlenmeyer flask and incubated at 30°C for 3 hours.

*Vahlkampfia avara* cells were maintained in cyst phase in static flasks containing freshwater base (FWB) medium (2.2 mM NaCl, 22 μM MgSO_4_, 34 μM CaCl_2_·2H_2_O, 1mM Na_2_HPO_4_, 1 mM KH_2_PO_4_) at 22°C, then grown on FW plates (freshwater base recipe and 1.5% agar) containing a lawn of *E. coli*.

*Dictyostelium discoideum* cells (strain AX2-ME, cloned and expanded from AX2-RRK DBS0235521 obtained from the Dictybase stock center^61^) were grown axenically in HL5 media (Formedium, HLE2) at 21 °C, and split into fresh media every 2-3 days through a ∼50-100 fold dilution. Cells were used for experiments between passage 3 and 20.

### *Naegleria* protein extraction

Lysis Buffer (100 mM Tris pH 8.1, 2.5 mM EGTA, 1 mM MgCl_2_, 0.5% TritonX-100, 1X HALT, 0.2 mM PMSF) and Gel Sample Buffer (10 mM DTT in Invitrogen NuPAGE 4X LDS Buffer solution) were prepared fresh just prior to use. Prior to extraction, *Naegleria* cells were cultured for 48 hours (to confluency) in T75 flasks containing 15 ml of M7 medium. Cells were washed once in 2 mM Tris before centrifuging at 850 RCF for 5 minutes at room temperature. The cell pellet was immediately resuspended in 100 μl of lysis buffer and gently agitated for 5 minutes. 50% of the volume (50 μl) was collected for “total cell lysate,” then combined with 50 μl of gel sample buffer. The remaining fraction was centrifuged in a pre-chilled Beckman Coulter benchtop centrifuge at 15000 RCF for 5 minutes at 4°C. The resulting supernatant was collected from the pellet, and each fraction was resuspended in 50 μl of gel sample buffer. All three fractions were heated for 3 minutes at 90°C before freezing at -80°C in a pre-chilled container.

### Custom Antibodies

Custom rabbit polyclonal antibodies were raised against the three *Naegleria gruberi* Myosin 2 proteins with synthetic peptides: Myo2A (GenBank: XP_002671980.1) using amino acids 2-30 (SQENDNIEVGDESQRRRRTARGGAGAGTK) and amino acids 1896-1915 (KTQRLARKASPFSASQLVDD) as peptide antigens, Myo2B (GenBank:XP_002680829.1) using amino acids 12-35 (KKSGGDNKSSFFQRLLEQQKSDQA) and amino acids 941-961 (DQAKRQLSDKDVEVLQRGKSV) as peptide antigens, and Myo2C (unannotated) using amino acids 2-22 (SRNRKSTNTEIIEDESQYKIK) and amino acids 1950-1972 (SGLLKEDDETVAADSSAPVEAEE) as peptide antigens. Peptide synthesis, antibody generation and affinity purification were performed by Pacific Immunology (Ramona, CA).

### SDS-PAGE

5 μl of each sample fraction was loaded into a 4-20% gradient BioRad TGX stain-free gel, along with a with Precision Plus protein standard (BioRad 161-0374) and electrophoresed in a BioRad (mini box) at 200V for 30 minutes. The gels were briefly rinsed in MilliQ water before they were activated with UV light for 5 minutes and imaged.

### Western blotting

PVDF membranes were rehydrated per manufacturer’s protocol, then equilibrated in Towbin’s transfer buffer without methanol (25 mM Tris, 192 mM glycine, pH 8.3). Samples were transferred from stain-free gels for 16 hours at 20V and 4C. The transfer was verified by imaging using UV, then blocked for 30 minutes in 5% milk-TBS-T (20 mM Tris pH 7.5, 137 mM NaCl, 0.5% (v/v) Tween-20) + 5% skim milk (Difco) at room temperature. Primary antibodies were diluted 1:200 in blocking solution and incubated at room temperature for 2 hours. The membrane was washed 3x in TBST for 5 minutes each followed by an additional 15 minute rinse. Membranes were then probed with secondary antibody (anti-rabbit conjugated to horseradish peroxidase, Thermo Fisher Cat# G-21234) diluted 1:10,000 in blocking solution, and incubated 1 hour at room temperature, then washed as before. Chemiluminescence substrate was prepared just prior to imaging following the manufacturer’s instructions (Amersham ECL Prime, Cat# RPN2232), and visualized on a GBox XX9 gel imager.

### Western blot quantitation

Antibody intensity peaks were measured for each lane in Fiji^62^ using the built-in gel analyzer function. Peaks were isolated by selecting each lane, plotting the band intensities, subtracting the background, then measuring the area under each peak. Total protein values were quantified from the corresponding lanes on the PVDF image taken after transfer. Western blot bands were then normalized by dividing each peak area by the total protein area.

### Immunofluorescence

Cells were collected and washed with 2 mM Tris pH 7.2, added to plasma-cleaned glass-bottomed 96-well plates coated with 0.1% fish skin gelatin and allowed to adhere for 5 minutes. Cells were fixed for 15 minutes by adding an equal volume of 2X PFA fixative (50 mM sodium phosphate buffer pH 7.2, 125 mM sucrose, and 3.6% paraformaldehyde). Cells were rinsed twice in PEM (100 mM PIPES, 1 mM EGTA, 0.1 mM MgSO_4_; pH 7.4) and permeabilized for 10 minutes in PEM + 0.05% TritonX-100 + 6.6 nM Alexa Fluor 488 Phalloidin (Invitrogen™ A12379). Cells were then blocked for one hour at room temperature in PEMBALG (PEM + 1% BSA, 0.1% sodium azide, 100 mM lysine, and 0.5% cold water fish skin gelatin; pH 7.4) followed by incubation for one hour at room temperature with primary antibodies (purified custom rabbit-anti-Myosin 2 antibodies generated by Pacific Immunology) diluted to 1-5 μg/ml in PEMBALG. Cells were then washed 3 times in PEMBALG and incubated at room temperature for one hour in secondary antibody (Alexa Fluor 555 conjugated goat anti-rabbit, (Invitrogen™ A21428) diluted to 2 μg/ml, with 6.6 nM Alexa Fluor 488 Phalloidin and 1 μg/ml DAPI. Cells were then rinsed 3X in PEM and imaged the same day.

### Ghost immunofluorescence

Cells were collected and washed as for immunofluorescence then allowed to adhere in 10 mg/ml PEI-coated coverslip-bottom 96-well plates for 5 minutes, then equilibrated 5 minutes in P buffer (10 mM PIPES, 10 mM EGTA, 1 mM MgCl_2_, 50 mM KCl, 1 mM DTT, pH 7.6). They were then extracted for 5 minutes with P buffer supplemented with 0.05% Triton X-100, 1 unit (1X) Halt™ Protease Inhibitor Cocktail, and 13.2 nM Alexa Fluor 488 Phalloidin, then washed twice with P buffer. An equal volume of 2X PFA fixative solution was immediately added to each well for 10 minutes at room temperature. They were then rinsed twice in PEM, treated as before for immunofluorescence, andimaged the same day without mounting media.

### Salt-treated ghost immunofluorescence

Cells were extracted as described above to create ghosts, followed by two washes in P-buffer. Each well was treated for 10 minutes with P buffer, P buffer supplemented with 200 mM KCl, P buffer with 1 mM Mg-ATP, or a sequential treatment of P buffer with KCl followed by Mg-ATP. All wells were then rinsed in P buffer, fixed, and stained with antibodies as described above. Each treatment was imaged the same day.

### ATP-induced ghost contraction

*Naegleria* NEG-M and *Vahlkampfia avara* cells were prepared on 5 mg/ml PEI in a 96 well plate and extracted, as described for immunofluorescence. The extracted ghosts were then washed twice in fresh P buffer and imaged during addition of either P buffer or P buffer containing 2 mM Mg-ATP (for a final concentration of 1 mM Mg-ATP). Images were taken at 500 millisecond intervals at a single z-slice just above the surface of the glass during addition of Mg-ATP. *Dictyostelium discoideum* cells were treated as described above with the following changes: cells in HL5 media were seeded into 24- or 96-well glass-bottom plates coated with 0.05 mg/mL PEI, and were then washed into P buffer. Images were taken at 3 second intervals at a single z-slice just above the surface of the glass during addition of Mg-ATP.

### Salt-treated ghost contraction

Ghosts were prepared as described above, washed twice in P buffer, then treated with either P buffer or P buffer containing 200 mM KCl for 10 minutes. The ghosts were rinsed again in P buffer, and imaged during addition of buffer containing Mg-ATP to a final concentration of 1 mM Mg-ATP as described above.

### Imaging

All imaging unless otherwise noted was done using a Nikon ECLIPSE Ti2 microscope equipped with a Plan Apo λ 100x oil objective (1.45 NA), a Crest spinning disk (50 μm), a Prime 95B 25mm CMOS camera (01-PRIME-95B-25MM-R-M-16-C), a Spectra III/Celesta Light Engine (P/N 90-10-512) for confocal illumination (at 50% power with excitation wavelengths of 405, 477, 546, and 638 nm) and a Sola light source (Lumencor, P/N 90-10444) for epifluorescence. The microscope was controlled through NIS Elements software. Fixed cell images were all acquired as multi-channel z stacks with a step size of 0.2 μm. *Dictyostelium* cells were imaged using a Nikon Ti2 microscope with a Plan Apo λ 20x air objective (0.75 NA), a Prime BSI Express camera, and a Sola II light source for widefield fluorescence.

### Live imaging and migration analysis

*Naegleria* and *Vahlkampfia* cells were resuspended in 2 mM Tris pH 7.2 and added to an untreated glass-bottom 96 well plate. Cells were allowed to adhere for >5 minutes, and imaged using DIC over 90 seconds with less than 1 second frame intervals. Cells were tracked in Fiji using the MTrackJ plugin^63^ and their trajectories, mean velocities, and directional persistence were reported for cells within the field of view at the beginning of the imaging session. Directional persistence of each cell was calculated by dividing the maximum displacement by the total path length, making a value of 1 a directly linear route.

### Colocalization analysis

Fluorescence channels were compared by manually selecting the indicated ROI, which was fully contained within the cell and avoided negative space where possible (vacuoles and nuclei) (**Dataset S1**). Co-localization between ROI for the two channels was then estimated using the built-in Fiji^62^ “colocalize” tool for the starting ROI, or with one channel rotated 90 degrees as a randomized pixel control. All colocalization measurements were made using Fay randomization over 25 iterations. Pixel-by-pixel signal covariance was normalized by their standard deviations from mean intensity. The resultant values ranged from 1.0 to -1.0, where 1.0 indicates perfect correlation, -1.0 indicates perfect anti-correlation, and values close to 0 indicated no correlation. Line scans were created using the “plot profile” tool as indicated and X,Y coordinates plotted in GraphPad Prism software. Lines were normalized to highest and lowest values and cropped to exclude regions outside of the cell.

### Image analyses; total fluorescence measurements

Maximum intensity projections of confocal images of immunostained ghosts were thresholded using the IsoData method in Fiji, and “Analyze Particles” tool was used on the actin channel to identify total ghost areas and generate masks for each ghost. The fluorescence intensity of the total surface area was measured for each channel within the total ghost area. These measurements were then used to calculate the mean fluorescence intensity for each Myo2 antibody and graphed using Prism.^64^

### Image analyses; calculating cell size for ghost contraction experiments

*Dictyostelium* ghost contraction experiments were quantified in Fiji^62^ using the magic wand and measure tools to determine the cell area before and <2 minutes after the addition of Mg-ATP or P buffer. For each of three separate experimental trials, 2 to 3 wells were used as replicates. 27 to 45 cells were analyzed per treatment condition per experiment, and were randomly selected from the field of view in the pretreatment image. If a cell left the focal plane during treatment, a different cell was selected when available. Data were plotted in GraphPad Prism. *Naegleria* and *Vahlkampfia* ghost contraction experiments were quantified in Fiji using the magic wand tool as described above, for >30 seconds before and <2 minutes after addition of Mg-ATP or P buffer. Each experimental replicate included 1 to 4 wells with a minimum of 8 cells measured per treatment. All cells of the selected field of view were measured unless they left the focal plane.

### SuperPlots

Data in Figures 4, 5, and 6 were plotted in GraphPad prism as SuperPlots^64^ with semi-transparent points indicating single measurements, and separate experimental replicates differentiated by color and shape of points. The average for each experimental replicate is overlayed in a larger symbol of the same shape and color. N indicates the number of times the experiment was independently repeated.

### Myosin 2 sequence collection

We identified potential heterolobosean Myosin 2 sequences using four approaches: <u>1. *Manual NCBI search:*</u> Potential Myosin 2 sequences were identified by iterative BLAST searches (blastp and tblastn) of the NCBI and EukProt databases using the BLOSUM45 matrix, against all three *Naegleria gruberi* Myosin 2 peptide sequences. The searches were filtered to exclude Opisthokont (taxid:33154) and amoebozoan sequences (taxid:554915). For species with unassembled or unannotated genomes, resultant partial-length results were extended by selecting 5kb genomic regions upstream and downstream of the top hit and using the ExPASy Translate tool to identify candidate reading frames and checked for Myosin 2 protein domains and motifs using InterproScan. Translated sequences of appropriate length (1500-2100 residues) were added to the putative heterolobosean Myosin 2 protein sequence collection. <u>2. *TCS proteome search:*</u> Potential Myosin 2 sequences were identified using the The Comparative Set (TCS; 196 species chosen based on BUSCO completion and phylogenetic importance) from the Eukprot proteome set.^48^ Due to overrepresentation of Metazoa species in this data set, along with the atypical Myosin II family expansion in this lineage, we omitted Metazoan proteomes from this search. To generate a search seed, we collected all Myosin II sequences from an existing pan-eukaryotic Myosin set,^23^ aligned these using MAFFT v7.505 with its L-INS-i algorithm and the BLOSUM30 scoring matrix, manually removed the hypervariable regions on the ends of the alignment, and then removed sequences with truncated motor domains. We built a hidden Markov model (HMM) profile from the remaining 38 sequences, then used this profile to search for additional Myosin II sequences in our TCS database using Hmmer v3.3.2.^65^ We collected the ten highest scoring matches from each species, aligned these with MAFFT as above, and inferred an exploratory tree using IQ-tree2 v2.3.1.^66^ We examined the resultant consensus tree and the HMM bit scores for genes in the branch containing all Myosin 2s in comparison to non-Myosin II branches. Based on this, we thresholded candidate genes to those with HMM bit scores >1400. <u>3. *Searching Pánek 2025*</u> *<u>transcriptomes:</u>* 16 heterolobosean transcriptome assemblies^49^ were downloaded, then translated into predicted proteins using TransDecoder v5.5.0.^67^ These were then scanned using the same Myosin 2 motor HMM profile generated for the TCS set using Hmmer v3.3.2^65^ on default settings. The candidate hits were thresholded at a more permissive bitscore of >1000 to allow more divergent sequences. <u>4. *Sequencing and searching the* Vahlkampfia *transcriptome*:</u> *Vahlkampfia avara* amoebae were washed in 2 mM Tris then pelleted. Total RNA was extracted from 2 x 10^6 cells in TRIzol reagent, and enriched for poly(A) mRNA transcripts. RNAseq was performed by Illumina, with 2x151 bp paired-ends totaling 236M reads. Reads were assembled into a transcriptome using Trinity v2.15.1 default settings,^68^ then translated into predicted proteins using TransDecoder v5.5.0 default settings.^67^ This protein database was scanned using the PFAM^69^ Myosin motor head HMM (PF00063) using Hmmer v3.3.2^65^ and all candidate sequences scoring better than the default gathering threshold were collected.

### Gene tree estimation

To place our Myosin 2 candidates on a pan-eukaryotic Myosin scaffold tree, we combined our candidate full length Myosin 2 sequences with other Myosin protein sequences from representative eukaryotic species in an existing data set.^6^ This combined set of sequences was processed to remove duplicates, transcript variants, and aberrant sequences likely to be from sequencing errors using CD-HIT v4.8.1^70^ and TreeShrink v1.3.9.^71^ The remaining genes were aligned using MAFFT as above, then manually trimmed to the motor domain. This alignment was used to infer a maximum-likelihood gene tree using IQ-Tree, using its ModelFinder algorithm to select the best model of evolution (Q.yeast+R7 chosen according to BIC) with 5000 ultrafast bootstrap tests. We visualized the tree using a custom R script and the packages ggplot2 v3.5.1^72^ and ggrtee v3.6.2^73^ (**Figure S2**).

### Myosin 2 sequence validation and feature prediction

We validated putative Myosin 2 sequences based on the predicted protein length, domain organization, and phylogenetic placement. We included *Dictyostelium* Myosin 2 (XP_637740.1) as a reference sequence, and included *Acanthamoeba castellani* (XP_004358280.1) and human Mhc10 (NP_005955.3) as comparison, with *Dictyostelium* Myo1A (XP_641363.1) and Myo1D (XP_643446.1) as outgroups. The protein sequences were aligned in Geneious Prime v. 2025.1.2 using ClustalO,^74^ then truncated to include the N-terminus through the second IQ motif. These expanded motor alignments were organized phylogenetically with a tree built in Geneious using RAxML v 8.2.12. To clarify sub-domain sequence similarity of known functional motifs, we used a conservative substitution matrix (BLOSUM62) to determine the pairwise positive scores between each of the heterolobosean Myosin 2 sequences to the reference sequence of *Dictyostelium* Myosin 2. We distinguished between regions that had sequence similarities over 80%, 30-80% and less than 30%. Some missing regions (labeled “404”) resulted from incomplete datasets, as some heterolobosean Myosin 2 sequences were not from full-length assemblies.

---

**Figure S1:**
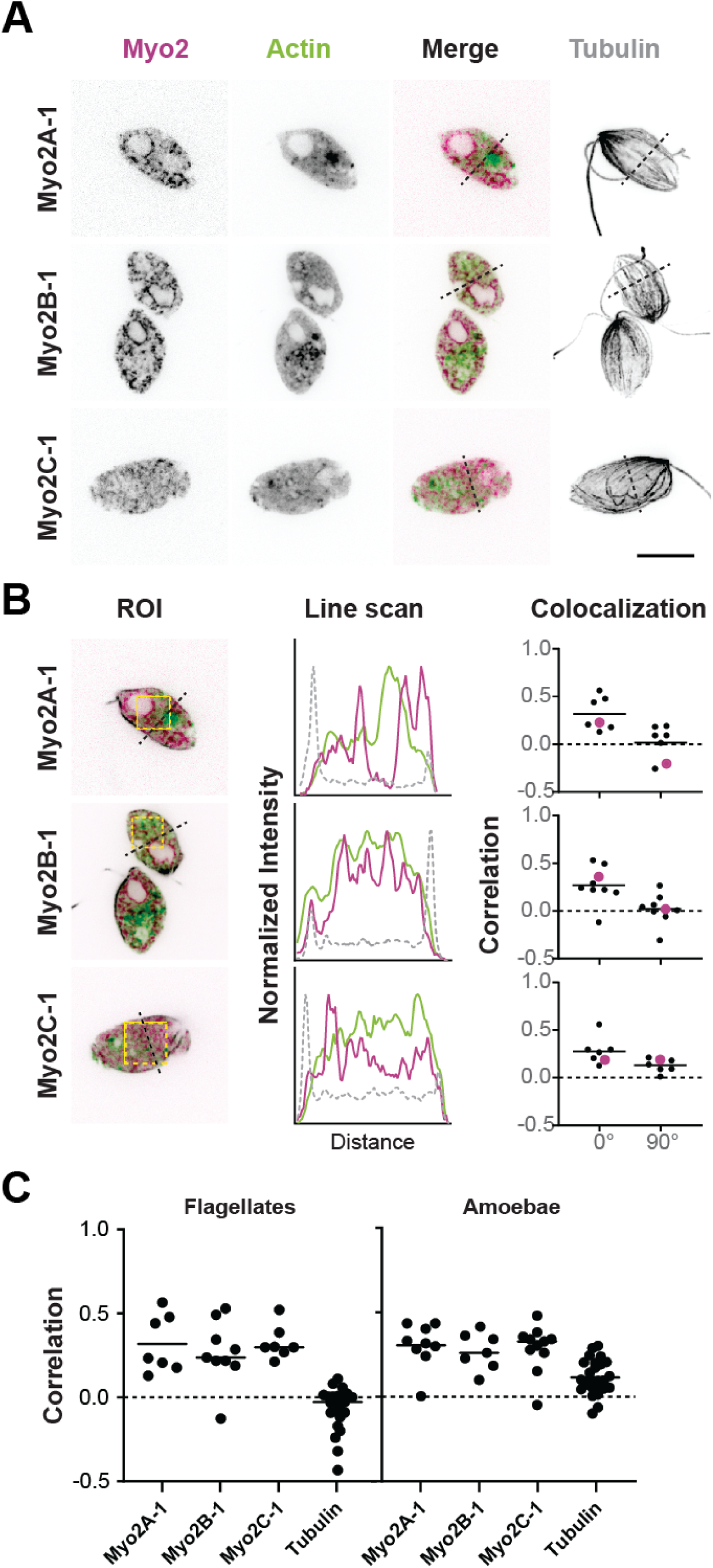
Myosin 2 localizes with actin in *Naegleria* flagellates. (**A**) *Naegleria* flagellates were fixed and stained with Myo2 antibodies (pink) to detect Myosin 2, DM1A to detect α-tubulin (gray), and Alexa Fluor™ 488 Phalloidin to label actin filaments (green). Single z slices through the midsection of cells are shown. (**B**) The Region of Interest (ROI) for each example depicts the area used for colocalization analyses; dotted yellow squares were used in 90° rotation tests, and dotted lines through the cell were used for line scans. The colocalization values are plotted as Pearson’s correlation coefficients, with the representative cells shown on the left indicated by pink circles. The Myo2 and actin line scan fluorescence intensity was normalized to the dimmest (0%) and brightest (100%) signal for regions within the cortex boundary. (n ≥ 6 cells for each antibody; scale bar = 10 μm). (**C**) A comparison of the Pearson’s correlation coefficients for Myosin 2:actin between flagellates and amoebae shows that the Myosin 2 signal localizes weakly (<0.5) to actin in both cell types, using tubulin:actin localization as a negative control.

**Figure S2:**
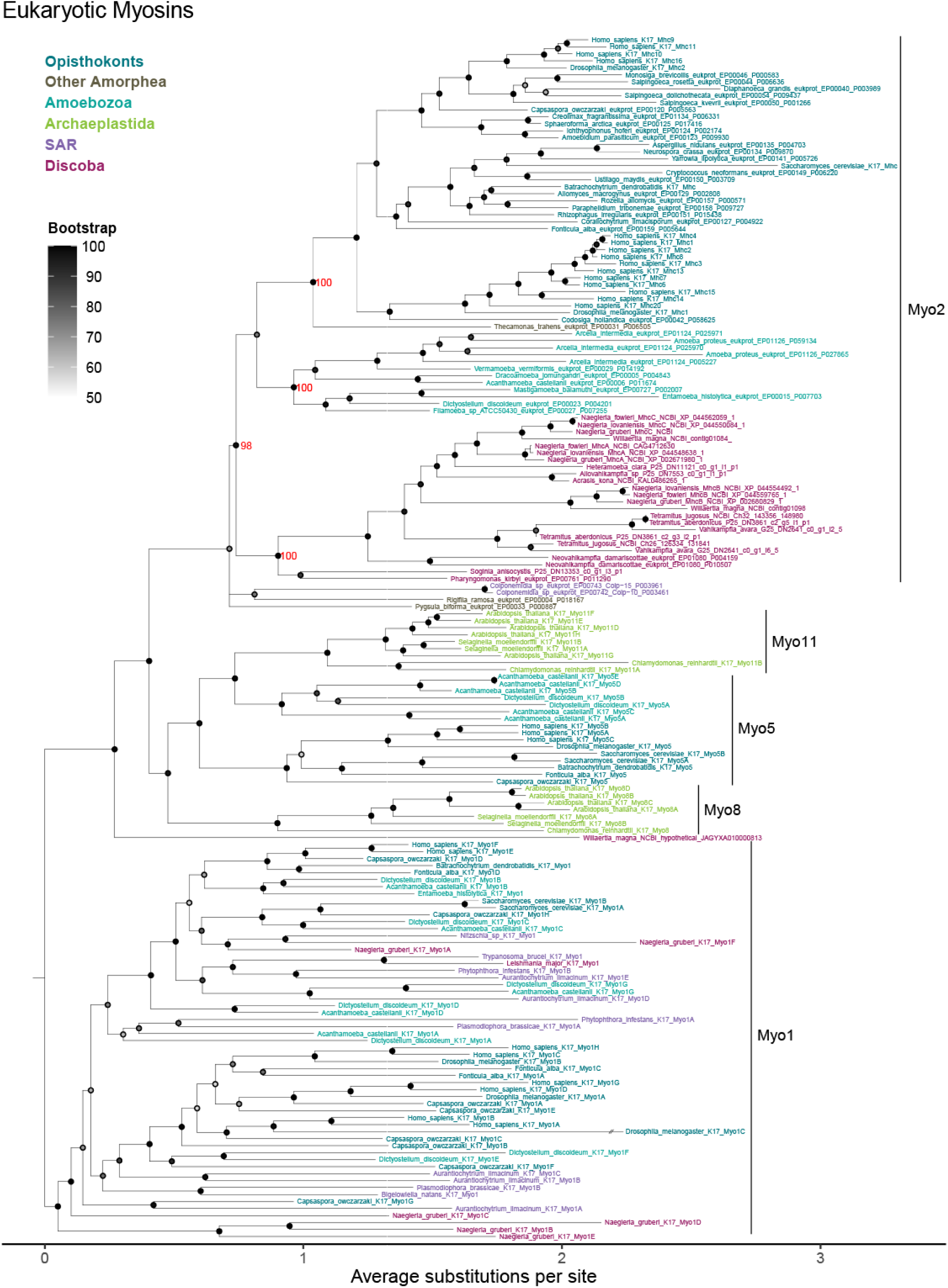
Heterolobosea encode direct orthologs to Amorphean Myosin 2s. Eukaryotic myosins were identified using an HMM built from Amorphean Myosin 2 sequences and are presented in a gene tree alongside other myosin types. The tree is rooted at the edge connecting the Myosin I branch to all other branches. Gene labels are color-coded by eukaryotic supergroup (upper left). Nodes with bootstrap support below 50% are collapsed into polytomies, and bootstrap values for key nodes are displayed in red. *Naegleria* and other heterolobosean Myosin 2s cluster with all other Myosin 2s (98% bootstrap support).

**Figure S3:**
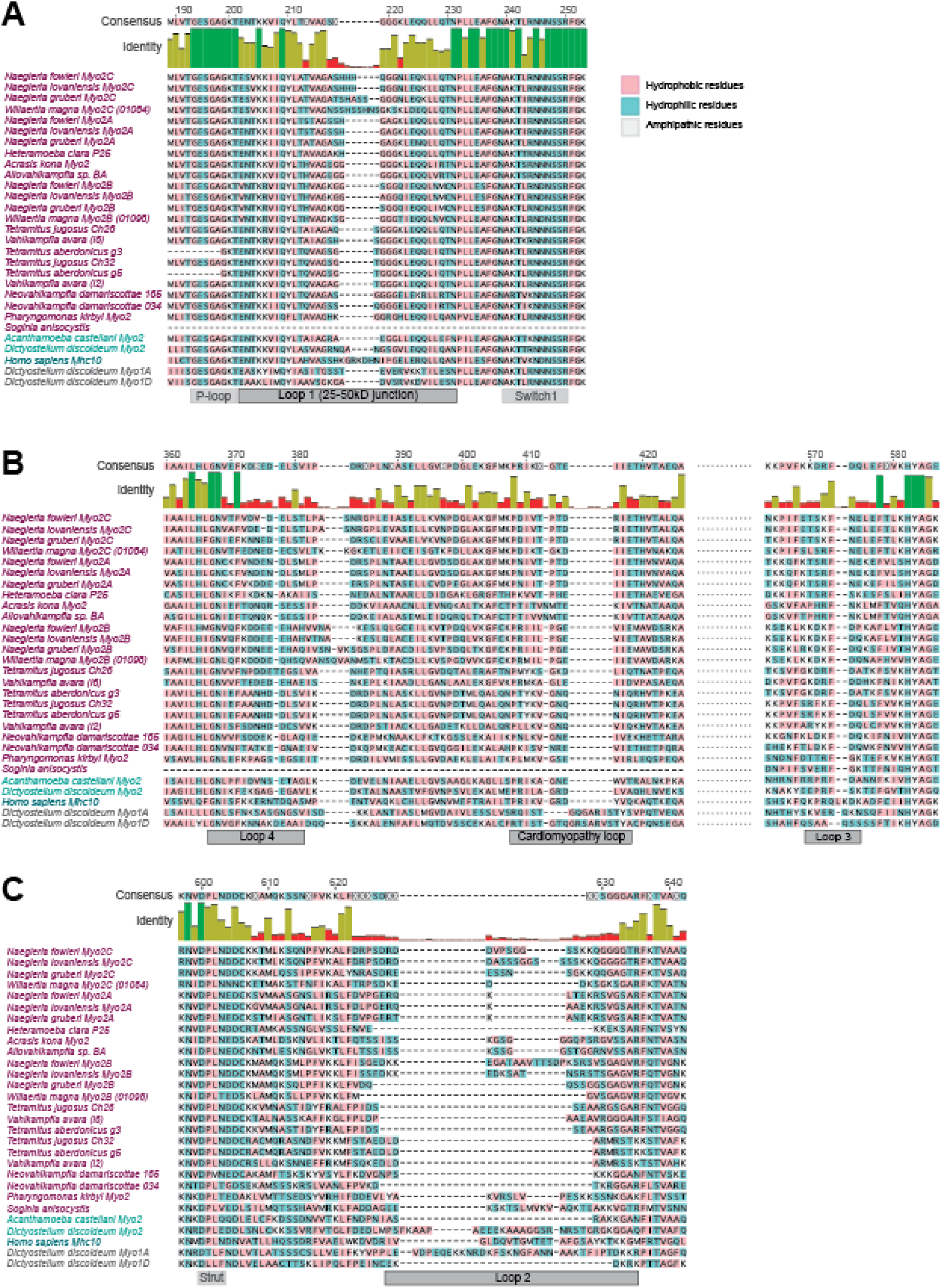
Heterolobosean Myosin 2 sequences vary from Amorphean Myosin 2s in their actin-binding regions. Myosin 2’s actin binding is dominated by strong hydrophobic interactions between surface residues on actin and corresponding residues of the motor domain’s actin-binding interface, including the ‘cardiomyopathy (CM) loop’ and loops 1-4.^55,56^ (**A-C**) A multi-sequence alignment using ClustalO shows heterolobosean sequence similarity of actin-binding regions: (**A**) Loop 1, (**B**) Loop 4, the cardiomyopathy (CM) loop, Loop 3 and (**C**) Loop 2. The residues are color-coded by hydrophobicity, with hydrophobic residues in pink, hydrophilic in blue, and amphipathic in gray. By pairwise comparison to *Dictyostelium*, the average heterolobosean sequence similarity was 70.2% (vs *Acanthamoeba* 75.8%) in Loop 1, 52.3% (vs *Acanthamoeba* 42.9%) in Loop 4, 51.9% (vs *Acanthamoeba* 63.6%) in CM loop, 59.4% (vs *Acanthamoeba* 50%) in Loop 3, and 28.0% (vs *Acanthamoeba* 44.4%) in Loop 2.

**Figure S4:**
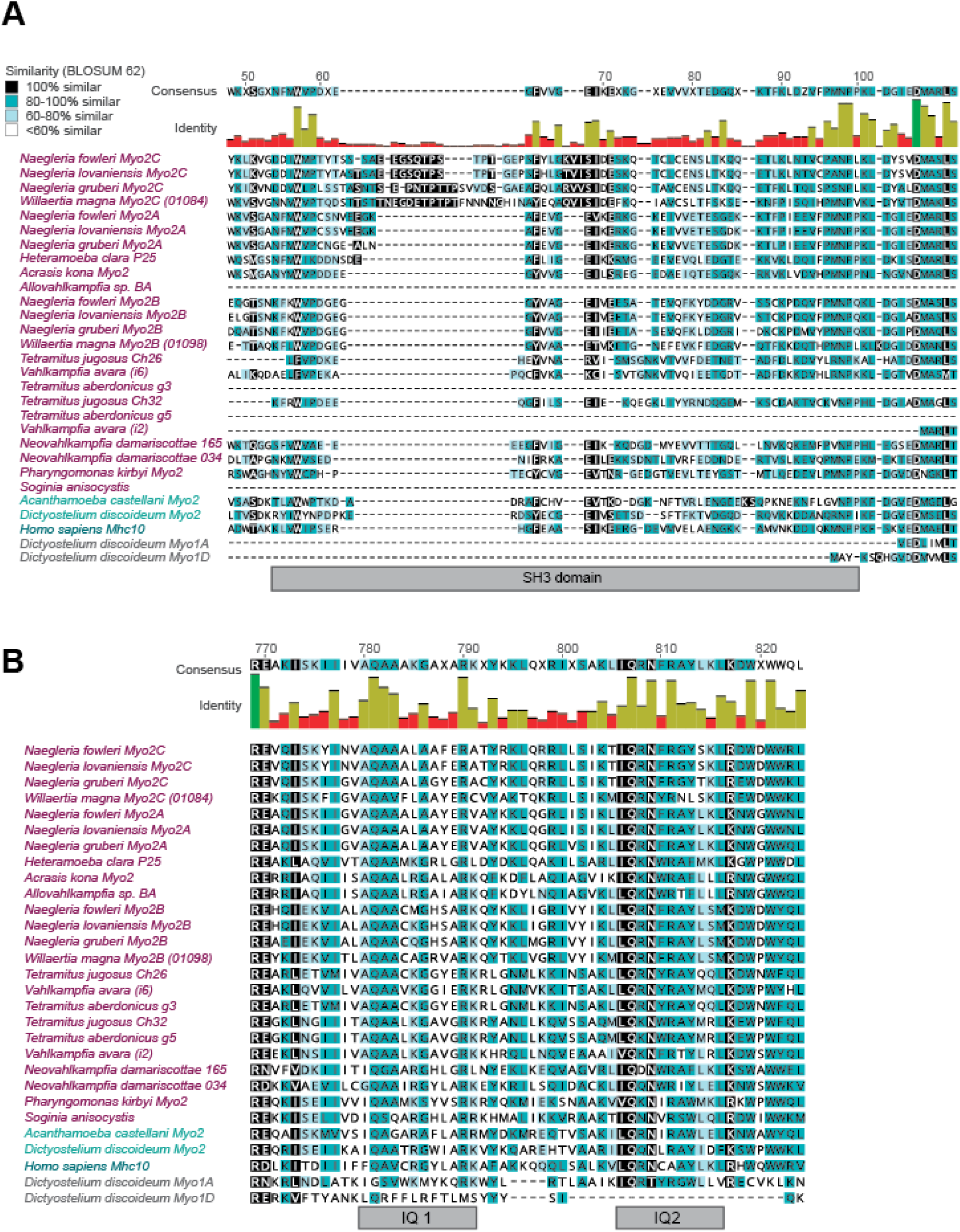
Similarity across regions outside the Myo2 motor domain. The SH3 domain at the N-terminus of Myosin 2 proteins contributes to its kinetic properties, actin affinity, and regulator interactions.^53,54,86^ (**A**) A multi-sequence alignment using ClustalO shows heterolobosean sequence similarity of the SH3 domain to the alignment consensus, colored by percent similarity (black = 100% BLOSUM62, teal = 80-100%, light blue = 60-80%, and white = <60%). The Heteroloboseans’ SH3 regions had variable scores with less than 80% similarity to the *Dictyostelium* Myo2 reference (average 31.5%, compared to *Acanthamoeba* at 36.4%). (**B**) The regulatory and essential light chains, which are critical to Myosin 2 function, interact with the two IQ motifs in the neck region beyond the motor domain.^57,87^ The IQ motifs in this alignment generally follow the IQ motif consensus. By pairwise comparison to the *Dictyostelium* Myosin 2 reference, the first IQ motif averaged 56.5% similar for Heteroloboseans, (vs *Acanthamoeba* 75%), and the second IQ motif was 67.6% similar for Heteroloboseans, (vs *Acanthamoeba* 90.9%).

## SUPPLEMENTAL DATASET AND MOVIE LEGENDS

**Dataset S1:** A spreadsheet with manually selected ROIs and colocalization analyses shown in **Figure 3A** and **Figure S1**. This spreadsheet has four tabs: Flagellate ROI, Amoeba ROI, Flagellate Colocalization, and Amoeba Colocalization. Rows in ROI tabs correspond to paired images of individual cells stained for Myosin 2, actin (green) and tubulin (gray). Each image shows the ROI (yellow box) used for colocalization measurements of both Myo2:actin and tubulin:actin. The Pearson’s correlation coefficients (R) were measured for each ROI at 0° and 90° rotations. In the Flagellate and Amoeba Colocalization tabs, the Pearson’s correlation coefficients are organized first by antibodies used (Myo2A-C and tubulin), then by degree of ROI rotation (0 and 90). Below these columns are the averages for each ROI’s R-value (“Average Pearson’s”), SE for all R-values, and the combined averages and SE for all 3 sets of both Myo2:actin and tubulin:actin, indicated by 0 or 90 degrees.

**Movie S1:** A video showing three example ghosts of *Dictyostelium, Naegleria,* and *Vahlkampfia* contracting upon addition of Mg-ATP. Each time series is labeled, and each series was adjusted by removing time points prior to treatment to show Mg-ATP addition at approximately the same time. All three ghosts are shown at the same scale (Time = minutes:seconds, scale bar = 10 μm).

## Notes

### Competing Interest Statement

The authors have declared no competing interest.

## REFERENCES

1. García-Arcos, J.M., Jha, A., Waterman, C.M., and Piel, M. (2024). Blebology: principles of bleb-based migration. Trends Cell Biol. 34, 838–853.

2. Svitkina, T. (2018). The Actin Cytoskeleton and Actin-Based Motility. Cold Spring Harb. Perspect. Biol. 10. 10.1101/cshperspect.a018267.

3. Weißenbruch, K., and Mayor, R. (2024). Actomyosin forces in cell migration: Moving beyond cell body retraction. Bioessays 46, e2400055.

4. Niederman, R., and Pollard, T.D. (1975). Human platelet myosin. II. In vitro assembly and structure of myosin filaments. J. Cell Biol. 67, 72–92.

5. Quintanilla, M.A., Hammer, J.A., and Beach, J.R. (2023). Non-muscle myosin 2 at a glance. J. Cell Sci. 136, jcs260890.

6. Kollmar, M., and Mühlhausen, S. (2017). Myosin repertoire expansion coincides with eukaryotic diversification in the Mesoproterozoic era. BMC Evol. Biol. 17, 211.

7. Adl, S.M., Simpson, A.G.B., Lane, C.E., Lukeš, J., Bass, D., Bowser, S.S., Brown, M.W., Burki, F., Dunthorn, M., Hampl, V., et al. (2012). The revised classification of eukaryotes. J. Eukaryot. Microbiol. 59, 429–493.

8. Vicente-Manzanares, M., Ma, X., Adelstein, R.S., and Horwitz, A.R. (2009). Non-muscle myosin II takes centre stage in cell adhesion and migration. Nat. Rev. Mol. Cell Biol. 10, 778–790.

9. Charras, G., and Paluch, E. (2008). Blebs lead the way: how to migrate without lamellipodia. Nat. Rev. Mol. Cell Biol. 9, 730–736.

10. Seveau de Noray, V., Manca, F., Mainil, I., Remson, A., Biarnes-Pelicot, M., Gabriele, S., Valignat, M.-P., and Theodoly, O. (2022). Keratocytes migrate against flow with a roly-poly-like mechanism. Proc. Natl. Acad. Sci. U. S. A. 119, e2210379119.

11. Svitkina, T.M., Verkhovsky, A.B., McQuade, K.M., and Borisy, G.G. (1997). Analysis of the actin-myosin II system in fish epidermal keratocytes: mechanism of cell body translocation. J. Cell Biol. 139, 397–415.

12. Blaser, H., Reichman-Fried, M., Castanon, I., Dumstrei, K., Marlow, F.L., Kawakami, K., Solnica-Krezel, L., Heisenberg, C.-P., and Raz, E. (2006). Migration of zebrafish primordial germ cells: a role for myosin contraction and cytoplasmic flow. Dev. Cell 11, 613–627.

13. Yoshida, K., and Soldati, T. (2006). Dissection of amoeboid movement into two mechanically distinct modes. J. Cell Sci. 119, 3833–3844.

14. Wessels, D., Soll, D.R., Knecht, D., Loomis, W.F., De Lozanne, A., and Spudich, J. (1988). Cell motility and chemotaxis in *Dictyostelium* amebae lacking myosin heavy chain. Dev. Biol. 128, 164–177.

15. Chang, F., Kong, S.J., Wang, L., Choi, B.K., Lee, H., Kim, C., Kim, J.M., and Park, K. (2020). Targeting actomyosin contractility suppresses malignant phenotypes of acute myeloid leukemia cells. Int. J. Mol. Sci. 21, 3460.

16. Richards, T.A., and Cavalier-Smith, T. (2005). Myosin domain evolution and the primary divergence of eukaryotes. Nature 436, 1113–1118.

17. Preston, T.M., and King, C.A. (1978). An experimental study of the interaction between the soil amoeba Naegleria gruberi and a glass substrate during amoeboid locomotion. J. Cell Sci. 34, 145–158.

18. Strassert, J.F.H., Irisarri, I., Williams, T.A., and Burki, F. (2021). A molecular timescale for eukaryote evolution with implications for the origin of red algal-derived plastids. Nat. Commun. 12, 1879.

19. Preston, T.M., and King, C.A. (2003). Locomotion and phenotypic transformation of the amoeboflagellate *Naegleria gruberi* at the water-air interface. J. Eukaryot. Microbiol. 50, 245–251.

20. Velle, K.B., and Fritz-Laylin, L.K. (2020). Conserved actin machinery drives microtubule-independent motility and phagocytosis in *Naegleria*. J. Cell Biol. 219. 10.1083/jcb.202007158.

21. Russell, J.J., Theriot, J.A., Sood, P., Marshall, W.F., Landweber, L.F., Fritz-Laylin, L., Polka, J.K., Oliferenko, S., Gerbich, T., Gladfelter, A., et al. (2017). Non-model model organisms. BMC Biol. 15, 55.

22. Fritz-Laylin, L.K., Prochnik, S.E., Ginger, M.L., Dacks, J.B., Carpenter, M.L., Field, M.C., Kuo, A., Paredez, A., Chapman, J., Pham, J., et al. (2010). The genome of *Naegleria gruberi* illuminates early eukaryotic versatility. Cell 140, 631–642.

23. Sebé-Pedrós, A., Grau-Bové, X., Richards, T.A., and Ruiz-Trillo, I. (2014). Evolution and classification of myosins, a paneukaryotic whole-genome approach. Genome Biol. Evol. 6, 290–305.

24. Kennard, A.S., Velle, K.B., Ranjan, R., Schulz, D., and Fritz-Laylin, L.K. (2025). Tubulin sequence divergence is associated with the use of distinct microtubule regulators. Curr. Biol. 35, 233–248.e8.

25. Velle, K.B., Kennard, A.S., Trupinić, M., Ivec, A., Swafford, A.J.M., Nolton, E., Rice, L.M., Tolić, I.M., Fritz-Laylin, L.K., and Wadsworth, P. (2022). *Naegleria’s* mitotic spindles are built from unique tubulins and highlight core spindle features. Curr. Biol. 10.1016/j.cub.2022.01.034.

26. Burgess, D.R. (2005). Cytokinesis: new roles for myosin. Curr. Biol. 15, R310–R311.

27. Lai, E.Y., Walsh, C., Wardell, D., and Fulton, C. (1979). Programmed appearance of translatable flagellar tubulin mRNA during cell differentiation in *Naegleria* . Cell 17, 867–878.

28. Fulton, C., and Dingle, A.D. (1971). Basal bodies, but not centrioles, in *Naegleria*. J. Cell Biol. 51, 826–836.

29. Walsh, C.J. (2007). The role of actin, actomyosin and microtubules in defining cell shape during the differentiation of *Naegleria* amebae into flagellates. Eur. J. Cell Biol. 86, 85–98.

30. Fritz-Laylin, L.K., Assaf, Z.J., Chen, S., and Cande, W.Z. (2010). *Naegleria gruberi* de novo basal body assembly occurs via stepwise incorporation of conserved proteins. Eukaryot. Cell 9, 860–865.

31. Dunn, K.W., Kamocka, M.M., and McDonald, J.H. (2011). A practical guide to evaluating colocalization in biological microscopy. Am. J. Physiol. Cell Physiol. 300, C723–C742.

32. Yumura, S. (1991). Contraction of *Dictyostelium* ghosts reconstituted with myosin II. Cell Struct. Funct. 16, 481–488.

33. Analysis of proteostasis during aging with western blot of detergent-soluble and insoluble protein fractions Star Protocols. https://star-protocols.cell.com/protocols/753.

34. Yumura, S., and Fukui, Y. (1985). Reversible cyclic AMP-dependent change in distribution of myosin thick filaments in *Dictyostelium*. Nature 314, 194–196.

35. Podolski, J.L., and Steck, T.L. (1990). Length distribution of F-actin in *Dictyostelium discoideum*. J. Biol. Chem. 265, 1312–1318.

36. Xu, X.S., Lee, E., Chen, T., Kuczmarski, E., Chisholm, R.L., and Knecht, D.A. (2001). During multicellular migration, myosin ii serves a structural role independent of its motor function. Dev. Biol. 232, 255–264.

37. Kolega, J. (1997). Asymmetry in the distribution of free versus cytoskeletal myosin II in locomoting microcapillary endothelial cells. Exp. Cell Res. 231, 66–82.

38. Kaminer, B., and Bell, A.L. (1966). Myosin filamentogenesis: effects of pH and ionic concentration. J. Mol. Biol. 20, 391–401.

39. Gurmessa, B.J., Rust, M.J., Das, M., Ross, J.L., and Robertson-Anderson, R.M. (2021). Salt-mediated stiffening, destruction, and resculpting of actomyosin network. Front. Phys. 9. 10.3389/fphy.2021.760340.

40. Truong, T., Medley, Q.G., and Côté, G.P. (1992). Actin-activated Mg-ATPase activity of *Dictyostelium* myosin II. Effects of filament formation and heavy chain phosphorylation. J. Biol. Chem. 267, 9767–9772.

41. O’Halloran, T.J., Ravid, S., and Spudich, J.A. (1990). Expression of *Dictyostelium* myosin tail segments in *Escherichia coli*: domains required for assembly and phosphorylation. J. Cell Biol. 110, 63–70.

42. Rüegg, C., Veigel, C., Molloy, J.E., Schmitz, S., Sparrow, J.C., and Fink, R.H.A. (2002). Molecular motors: force and movement generated by single myosin II molecules. News Physiol. Sci. 17, 213–218.

43. Reines, D., and Clarke, M. (1985). Immunochemical analysis of the supramolecular structure of myosin in contractile cytoskeletons of *Dictyostelium* amoebae. J. Biol. Chem. 260, 14248–14254.

44. Sheikh, S., Fu, C.-J., Brown, M.W., and Baldauf, S.L. (2024). The *Acrasis kona* genome and developmental transcriptomes reveal deep origins of eukaryotic multicellular pathways. Nat. Commun. 15, 10197.

45. Liechti, N., Schürch, N., Bruggmann, R., and Wittwer, M. (2018). The genome of *Naegleria lovaniensis*, the basis for a comparative approach to unravel pathogenicity factors of the human pathogenic amoeba *N. fowleri*. BMC Genomics 19, 654.

46. Green, D.H., Rad-Menéndez, C., Culture Collection of Algae and Protozoa collective, Earlham Institute Genome Acquisition Lab and Protists Project, University of Oxford and Wytham Woods Genome Acquisition Lab, Darwin Tree of Life Barcoding collective, Wellcome Sanger Institute Tree of Life programme, Wellcome Sanger Institute Scientific Operations: DNA Pipelines collective, Tree of Life Core Informatics collective, and Darwin Tree of Life Consortium (2023). The genome sequence of the Heterolobosean amoeboflagellate, *Tetramitus jugosus* CCAP 1588/3C. Wellcome Open Res. *8*, 513.

47. Hasni, I., Chelkha, N., Baptiste, E., Mameri, M.R., Lachuer, J., Plasson, F., Colson, P., and La Scola, B. (2019). Investigation of potential pathogenicity of *Willaertia magna* by investigating the transfer of bacteria pathogenicity genes into its genome. Sci. Rep. 9, 18318.

48. Richter, D.J., Berney, C., Strassert, J.F.H., Poh, Y.-P., Herman, E.K., Muñoz-Gómez, S.A., Wideman, J.G., Burki, F., and de Vargas, C. (2022). EukProt: A database of genome-scale predicted proteins across the diversity of eukaryotes. Peer Community J. 2, article no. e56.

49. Pánek, T., Tice, A.K., Corre, P., Hrubá, P., Žihala, D., Kamikawa, R., Yazaki, E., Shiratori, T., Kume, K., Hashimoto, T., et al. (2025). An expanded phylogenomic analysis of Heterolobosea reveals the deep relationships, non-canonical genetic codes, and cryptic flagellate stages in the group. Mol. Phylogenet. Evol. 204, 108289.

50. Harding, T., Brown, M.W., Plotnikov, A., Selivanova, E., Park, J.S., Gunderson, J.H., Baumgartner, M., Silberman, J.D., Roger, A.J., and Simpson, A.G.B. (2013). Amoeba stages in the deepest branching heteroloboseans, including *Pharyngomonas*: evolutionary and systematic implications. Protist 164, 272–286.

51. Pánek, T., Simpson, A.G.B., Brown, M.W., and Dexter Dyer, B. (2016). Heterolobosea. In Handbook of the Protists (Springer International Publishing), pp. 1–42.

52. Tikhonenkov, D.V., Jhin, S.H., Eglit, Y., Miller, K., Plotnikov, A., Simpson, A.G.B., and Park, J.S. (2019). Ecological and evolutionary patterns in the enigmatic protist genus *Percolomonas* (Heterolobosea; Discoba) from diverse habitats. PLoS One 14, e0216188.

53. Fujita-Becker, S., Tsiavaliaris, G., Ohkura, R., Shimada, T., Manstein, D.J., and Sutoh, K. (2006). Functional characterization of the N-terminal region of myosin-2. J. Biol. Chem. 281, 36102–36109.

54. Lowey, S., Saraswat, L.D., Liu, H., Volkmann, N., and Hanein, D. (2007). Evidence for an interaction between the SH3 domain and the N-terminal extension of the essential light chain in class II myosins. J. Mol. Biol. 371, 902–913.

55. Sasaki, N., Ohkura, R., and Sutoh, K. (2003). *Dictyostelium* myosin II mutations that uncouple the converter swing and ATP hydrolysis cycle. Biochemistry 42, 90–95.

56. Sasaki, N., Asukagawa, H., Yasuda, R., Hiratsuka, T., and Sutoh, K. (1999). Deletion of the myopathy loop of *Dictyostelium* myosin II and its impact on motor functions. J. Biol. Chem. 274, 37840–37844.

57. Terrak, M., Wu, G., Stafford, W.F., Lu, R.C., and Dominguez, R. (2003). Two distinct myosin light chain structures are induced by specific variations within the bound IQ motifs-functional implications. EMBO J. 22, 362–371.

58. Pánek, T., Silberman, J.D., Yubuki, N., Leander, B.S., and Cepicka, I. (2012). Diversity, evolution and molecular systematics of the Psalteriomonadidae, the main lineage of anaerobic/microaerophilic heteroloboseans (excavata: discoba). Protist 163, 807–831.

59. Siddiqui, R., Ali, I.K.M., Cope, J.R., and Khan, N.A. (2016). Biology and pathogenesis of *Naegleria fowleri*. Acta Trop. 164, 375–394.

60. Cline, M., Carchman, R., and Marciano-Cabral, F. (1986). Movement of *Naegleria fowleri* stimulated by mammalian cells in vitro. J. Protozool. 33, 10–13.

61. Fey, P., Dodson, R.J., Basu, S., and Chisholm, R.L. (2013). One stop shop for everything *Dictyostelium*: dictyBase and the Dicty Stock Center in 2012. Methods Mol. Biol. 983, 59–92.

62. Schindelin, J., Arganda-Carreras, I., Frise, E., Kaynig, V., Longair, M., Pietzsch, T., Preibisch, S., Rueden, C., Saalfeld, S., Schmid, B., et al. (2012). Fiji: an open-source platform for biological-image analysis. Nat. Methods *9*, 676–682.

63. Meijering, E., Dzyubachyk, O., and Smal, I. (2012). Methods for cell and particle tracking. Methods Enzymol. 504, 183–200.

64. Lord, S.J., Velle, K.B., Mullins, R.D., and Fritz-Laylin, L.K. (2020). SuperPlots: Communicating reproducibility and variability in cell biology. J. Cell Biol. 219. 10.1083/jcb.202001064.

65. Eddy, S.R. (2011). Accelerated profile HMM searches. PLoS Comput. Biol. 7, e1002195.

66. Nguyen, L.-T., Schmidt, H.A., von Haeseler, A., and Minh, B.Q. (2015). IQ-TREE: a fast and effective stochastic algorithm for estimating maximum-likelihood phylogenies. Mol. Biol. Evol. 32, 268–274.

67. Haas, B.J. (2023). TransDecoder: TransDecoder source (Github).

68. Grabherr, M.G., Haas, B.J., Yassour, M., Levin, J.Z., Thompson, D.A., Amit, I., Adiconis, X., Fan, L., Raychowdhury, R., Zeng, Q., et al. (2011). Full-length transcriptome assembly from RNA-Seq data without a reference genome. Nat. Biotechnol. 29, 644–652.

69. El-Gebali, S., Mistry, J., Bateman, A., Eddy, S.R., Luciani, A., Potter, S.C., Qureshi, M., Richardson, L.J., Salazar, G.A., Smart, A., et al. (2019). The Pfam protein families database in 2019. Nucleic Acids Res. 47, D427–D432.

70. Li, W., and Godzik, A. (2006). Cd-hit: a fast program for clustering and comparing large sets of protein or nucleotide sequences. Bioinformatics 22, 1658–1659.

71. Mai, U., and Mirarab, S. (2018). TreeShrink: fast and accurate detection of outlier long branches in collections of phylogenetic trees. BMC Genomics 19, 272.

72. Wickham, H. (2016). Ggplot2 2nd ed. (Springer International Publishing).

73. Yu, G., Smith, D.K., Zhu, H., Guan, Y., and Lam, T.T.-Y. (2017). ggtree : an r package for visualization and annotation of phylogenetic trees with their covariates and other associated data. Methods Ecol. Evol. 8, 28–36.

74. Sievers, F., Wilm, A., Dineen, D., Gibson, T.J., Karplus, K., Li, W., Lopez, R., McWilliam, H., Remmert, M., Söding, J., et al. (2011). Fast, scalable generation of high-quality protein multiple sequence alignments using Clustal Omega. Mol. Syst. Biol. 7, 539.

75. Krishnan, D., and Ghosh, S.K. (2020). Morphological and motility features of the stable bleb-driven monopodial form of *Entamoeba* and its importance in encystation. Infect. Immun. 88. 10.1128/IAI.00903-19.

76. Cooper, M.S., and Schliwa, M. (1986). Motility of cultured fish epidermal cells in the presence and absence of direct current electric fields. J. Cell Biol. 102, 1384–1399.

77. Lei, X., Chen, X., Chen, J., and Liang, C. (2023). A new *Mayorella* species isolated from the Mariana Trench area (Pacific Ocean). Protist 174, 125958.

78. Toret, C., Picco, A., Boiero-Sanders, M., Michelot, A., and Kaksonen, M. (2022). The cellular slime mold *Fonticula alba* forms a dynamic, multicellular collective while feeding on bacteria. Curr. Biol. 32, 1961–1973.e4.

79. Zuppinger, C., and Roos, U.-P. (1997). Cell shape, motility and distribution of F-actin in amoebae of the mycetozoans *Protostelium mycophaga* and *Acrasis rosea*. A comparison with *Dictyostelium discoideum*. Eur. J. Protistol. 33, 396–408.

80. Siddiqui, R., and Khan, N.A. (2012). Biology and pathogenesis of *Acanthamoeba*. Parasit. Vectors 5, 6.

81. Táborský, P., Pánek, T., and Čepička, I. (2017). Anaeramoebidae fam. nov., a Novel Lineage of Anaerobic Amoebae and Amoeboflagellates of Uncertain Phylogenetic Position. Protist 168, 495–526.

82. Kusdian, G., Woehle, C., Martin, W.F., and Gould, S.B. (2013). The actin-based machinery of Trichomonas vaginalis mediates flagellate-amoeboid transition and migration across host tissue. Cell. Microbiol. 15, 1707–1721.

83. Fritz-Laylin, L.K., Lord, S.J., and Mullins, R.D. (2017). WASP and SCAR are evolutionarily conserved in actin-filled pseudopod-based motility. J. Cell Biol. 216, 1673–1688.

84. Parra-Acero, H., Harcet, M., Sánchez-Pons, N., Casacuberta, E., Brown, N.H., Dudin, O., and Ruiz-Trillo, I. (2020). Integrin-Mediated Adhesion in the Unicellular Holozoan Capsaspora owczarzaki. Curr. Biol. 30, 4270–4275.e4.

85. Gooday, A.J., Holzmann, M., Goetz, E., Cedhagen, T., Korsun, S., and Pawlowski, J. (2021). Three new species of *Gromia* (Protista, Rhizaria) from western Greenland fjords. Polar Biol. 44, 1037–1053.

86. Schröder, R.R., Manstein, D.J., Jahn, W., Holden, H., Rayment, I., Holmes, K.C., and Spudich, J.A. (1993). Three-dimensional atomic model of F-actin decorated with *Dictyostelium* myosin S1. Nature 364, 171–174.

87. Kollmar, M. (2006). Thirteen is enough: the myosins of *Dictyostelium discoideum* and their light chains. BMC Genomics 7, 183.

